# Comprehensive fitness maps of Hsp90 show widespread environmental dependence

**DOI:** 10.1101/823468

**Authors:** Julia M. Flynn, Ammeret Rossouw, Pamela A. Cote-Hammarlof, Ines Fragata, David Mavor, Carl Hollins, Claudia Bank, Daniel N.A. Bolon

## Abstract

Gene-environment interactions have long been theorized to influence molecular evolution. However, the environmental dependence of most mutations remains unknown. Using deep mutational scanning, we engineered yeast with all 44,604 single codon changes encoding 14,160 amino acid variants in Hsp90 and quantified growth effects under standard conditions and under five stress conditions. To our knowledge these are the largest determined comprehensive fitness maps of point mutants. The growth of many variants differed between conditions, indicating that environment can have a large impact on Hsp90 evolution. Multiple variants provided growth advantages under individual conditions, however these variants tended to exhibit growth defects in other environments. The diversity of Hsp90 sequences observed in extant eukaryotes preferentially contains variants that supported robust growth under all tested conditions. Rather than favoring substitutions in individual conditions, the long-term selective pressure on Hsp90 may have been that of fluctuating environments, leading to robustness under a variety of conditions.

## INTRODUCTION

The role of environment has been contemplated in theories of evolution for over a hundred years (Darwin, 1859; Darwin & Wallace, 1858; Wright, 1932), yet molecular level analyses of how environment impacts the evolution of gene sequences remain experimentally under-explored. Depending on environmental conditions, mutations can be categorized into three classes: deleterious mutations that are purged from populations by purifying selection, nearly-neutral mutations that are governed by stochastic processes, and beneficial mutations that provide a selective advantage (Ohta, 1973). It has long been clear that environmental conditions can alter the fitness effects of mutations (Tutt, 1896). However, examining how environmental conditions impact any of the three classes of mutations is challenging. Measurable properties of nearly-neutral and deleterious mutations in natural populations are impacted by both demography and selection (Ohta, 1973), which are difficult to disentangle. In addition, many traits are complex, making it challenging to identify all contributing genetic variations (McCarthy et al, 2008). For these and other reasons, we do not have a detailed understanding of how environmental conditions impact the evolution of most gene sequences.

Mutational scanning approaches (Fowler et al, 2010) provide novel opportunities to examine fitness effects of the same mutations under different laboratory conditions (Boucher et al, 2016; Boucher et al, 2014; Canale et al, 2018; Kemble et al, 2019). The EMPIRIC (Exceedingly Meticulous and Parallel Investigation of Randomized Individual Codons) approach that we developed is particularly well suited to address questions regarding the environmental impact of mutational effects for three reasons: it quantifies growth rates that are a direct measure of experimental fitness, all point mutations are engineered providing comprehensive maps of growth effects, and all the variants can be tracked in the same flask while experiencing identical growth conditions (Hietpas et al, 2011). We have previously used the EMPIRIC approach to investigate how protein fitness maps of ubiquitin vary in different environmental conditions (Mavor et al, 2016). The analysis of ubiquitin fitness maps revealed that stress environments can exacerbate the fitness defects of mutations. However, the small size of ubiquitin and the near absence of natural variation in ubiquitin sequences (only three amino acid differences between yeast and human) hindered investigation of the properties underlying historically observed substitutions.

Mutational scanning approaches have emerged as a robust method to analyze relationships between gene sequence and function, including aspects of environmentally dependent selection pressure. Multiple studies have investigated resistance mutations that enhance growth in drug or antibody environments (Dingens et al, 2019; Doud et al, 2018; Firnberg et al, 2014; Jiang et al, 2016; Stiffler et al, 2015). Most of these studies have focused on interpreting adaptation in the light of protein structure. Of note, Dandage, Chakraborty and colleagues explored how environmental perturbations to protein folding influenced tolerance of mutations in the 178 amino acid gentamicin-resistant gene in bacteria (Dandage et al, 2018). However, the question of how environmental variation shapes the selection pressure on gene sequences has not been well studied.

Here, we report comprehensive experimental fitness maps of Heat Shock Protein 90 (Hsp90) under multiple stress conditions and compare our experimental results with the historical record of hundreds of Hsp90 substitutions accrued during its billion years of evolution in eukaryotes. Hsp90 encodes a 709 amino acid protein and to our knowledge it is the largest gene for which a comprehensive protein fitness map has been determined. Hsp90 is an essential and highly abundant molecular chaperone which is induced by a wide variety of environmental stresses (Gasch et al, 2000; Lindquist, 1981). Hsp90 assists cells in responding to these stressful conditions by facilitating the folding and activation of client proteins through a series of ATP-dependent conformational changes mediated by co-chaperones (Krukenberg et al, 2011). These clients are primarily signal transduction proteins, highly enriched in kinases and transcription factors (Taipale et al, 2012). Through its clients, Hsp90 activity is linked to virtually every cellular process.

Hsp90 can facilitate the emergence and evolution of new traits in response to stress conditions, including drug resistance in fungi (Cowen & Lindquist, 2005), gross morphology in flies (Rutherford & Lindquist, 1998) and plants (Queitsch et al, 2002), and vision loss in cave fish (Rohner et al, 2013). In non-stress conditions, an abundance of Hsp90 promotes standing variation by masking the phenotypic effects of destabilizing mutations in clients. Stressful conditions that tax Hsp90 capacity can then manifest in phenotypic diversity that can contribute to adaptation. Because of the biochemical and evolutionary links between Hsp90 and stress, we hypothesized that environmental stress would result in altered fitness maps.

The conditions in natural environments often fluctuate, and all organisms contain stress response systems that aid in acclimation to new conditions. The conditions experienced by different populations can vary tremendously depending on the niches that they inhabit, providing the potential for distinct selective pressures on Hsp90. Previous studies of a nine amino acid loop in Hsp90 identified multiple amino acid changes that increased the growth rate of yeast in elevated salinity (Hietpas et al, 2013), demonstrating the potential for beneficial mutations in Hsp90. However, the sequence of Hsp90 is strongly conserved in eukaryotes (57% amino acid identity from yeast to human), indicating consistent strong purifying selection.

To investigate the potential influence of the environment on Hsp90 evolution, we quantified fitness maps in six different conditions. The different conditions impose distinct molecular constraints on Hsp90 sequence. While proximity to ATP is the dominant functional constraint in standard conditions, the influence of client and co-chaperone interactions on growth rate dramatically increases under stress conditions. Increased selection pressure from heat and diamide stresses led to a greater number of beneficial variants compared to standard conditions. The observed beneficial variants were enriched at functional hotspots in Hsp90. However, the natural variants of Hsp90 tend to support efficient growth in all environments tested, indicating selection for robustness to diverse stress conditions in the natural evolution of Hsp90.

## RESULTS

We developed a powerful experimental system to analyze the growth rate supported by all possible Hsp90 point mutations under distinct growth conditions. Bulk competitions of yeast with a deep sequencing readout enabled the simultaneous quantification of 98% of possible amino acid changes (Figure 1A). The single point mutant library was engineered by incorporating a single degenerate codon (NNN) into an otherwise wildtype Hsp90 sequence as previously described (Hietpas et al, 2012). To provide a sensitive readout of changes in Hsp90 function, we used a plasmid system that reduced Hsp90 protein levels to near-critical levels (Jiang et al, 2013). We employed a barcoding approach to efficiently track all variants in a single competition flask so that all variants experience identical conditions. As described in the Methods, the barcode strategy enabled us to track mutations across a large gene using a short sequencing readout. The barcoding strategy also reduced the impact of misreads as they result in unused barcodes that were discarded from the analyses.

**Figure 1.**
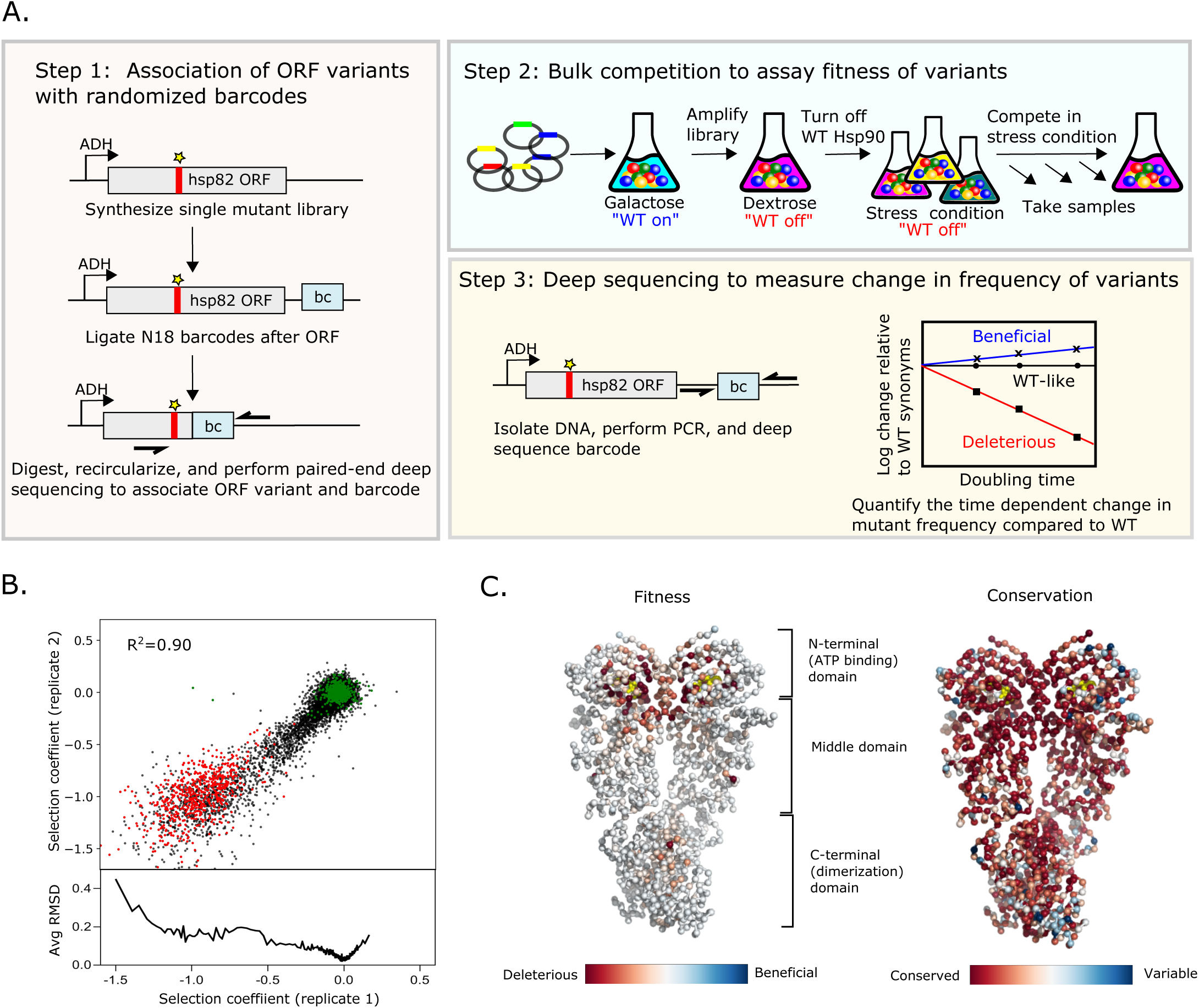
Approach to determine protein fitness maps of Hsp90. A. Barcoded competition strategy to analyze the growth effects of all single codon variants of Hsp90 in a single bulk culture. B. Measurements of selection coeffcients of all amino acid variants are reproducible in replicate growth competitions. Wildtype amino acids are shown in green and stop codons are shown in red. C. Average selection coeffcients at each position in standard conditions mapped onto a homodimeric structure of Hsp90 (Ali et al., 2006) and compared to patierns of evolutionary conservation. ATP is shown in yellow.

We transformed the plasmid library of comprehensive Hsp90 point mutations into a conditional yeast strain where we could turn selection of the library on or off. We used a yeast Hsp90-shutoff strain in which expression of the only genomic copy of Hsp90 is strictly regulated with a galactose-inducible promoter (Jiang et al, 2013). Yeast containing the mutant libraries were amplified under conditions that select for the plasmid, but not for the function of Hsp90 variants. We switched the yeast to dextrose media to shut off the expression of wildtype Hsp90 and then split the culture into six different environmental conditions. We extracted samples from each condition at multiple time points and used Illumina sequencing to estimate the frequency of each Hsp90 variant over time. We assessed the selection coefficient of each Hsp90 variant from the change in frequency relative to wildtype Hsp90 using a previously developed Bayesian MCMC method (Bank et al, 2014; Fragata et al, 2019).

To analyze reproducibility of the growth competition, we performed a technical replicate under standard conditions. We used a batch of the same transformed cells that we had frozen and stored such that the repeat bulk competition experiments and sequencing were performed independently. Selection coefficients between replicates were strongly correlated (R^2^=0.90), and indicated that we could clearly distinguish wildtype-like mutants from highly deleterious stop-like mutants (Figure 1B). The selection coefficients in this study also correlated strongly (R^2^=0.87) with estimates of the Hsp90 N-domain in a previous study (Mishra et al, 2016) (Figure S1A), indicating that biological replicates also show high reproducibility. Of note, variants with strongly deleterious effects exhibited the greatest variation between replicates, consistent with the noise inherent in estimating the frequency of rapidly depleting variants.

The large number of signaling pathways that depend on Hsp90 (Taipale et al, 2012) and its strong sequence conservation suggest that Hsp90 may be sensitive to mutation. However, most variants of Hsp90 were experimentally tolerated in standard conditions (Figure 1C, Figure S1B, and Table S1). All possible mutations were compatible with function at 425 positions. Only 18 positions had low mutational tolerance to the extent that 15 or more substitutions caused null-like growth defects (R32, E33, N37, D40, D79, G81, G94, I96, A97, S99, G118, G121, G123, Y125, F156, W300, and R380). All of these positions except for W300 are in contact with ATP or mediate ATP-dependent conformational changes in the N-domain of Hsp90. In fact, the average selection coefficient at different positions (a measure of mutational sensitivity) in standard growth conditions correlates (R^2^=0.49) with distance from ATP (Figure S1C). While W300 does not contact ATP, it transmits information from client binding to long range conformational changes of Hsp90 that are driven by ATP hydrolysis (Rohl et al, 2013). Our results indicate that ATP binding and the conformational changes driven by ATP hydrolysis impose dominant physical constraints in Hsp90 under standard laboratory conditions.

At first sight, the observation that most mutations are compatible with robust growth in standard conditions is at odds with the fact that the Hsp90 sequence is strongly conserved across large evolutionary distances (Figure 1C). One potential reason for this discrepancy could be that the strength of purifying selection in large natural populations over long evolutionary time-scales is more stringent than can be measured in the laboratory. In other words, experimentally unmeasurable fitness defects could be subject to purifying selection in nature. In addition, the range of environmental conditions that yeast experience in natural settings may not be reflected by standard laboratory growth conditions. To investigate the impact of environmental conditions on mutational effects in Hsp90, we measured the growth rate of Hsp90 variants under five additional stress conditions.

### Impact of stress conditions on mutational sensitivity of Hsp90

We measured the fitness of Hsp90 variants in conditions of nitrogen depletion (ND) (0.0125% ammonium sulfate), hyper-osmotic shock (0.8 M NaCl), ethanol stress (7.5% ethanol), the sulfhydryl-oxidizing agent diamide (0.85 mM), and temperature shock (37°C). All of these stresses are known to elicit a common shared environmental stress response characterized by altered expression of ∼900 genes as well as having specific responses unique to each stress (Gasch et al, 2000). Genes encoding heat shock proteins, including Hsp90, are transiently upregulated in all these stresses except elevated salinity (Gasch et al, 2000; Piper, 1995).

One way to characterize stress conditions is to measure the extent to which they slow down growth. For our experiments, each of the environmental stresses were selected to partially decrease the growth rate. Consistently, all stresses reduced the growth rate of the parental strain within a two-fold range, with depletion of nitrogen levels causing the smallest reduction in growth rate and diamide causing the greatest reduction (Figure 2A). To investigate how critical Hsp90 is for growth in each condition, we measured growth rates of yeast with either normal or more than 10-fold reduced (Jiang et al, 2013) levels of Hsp90 protein (Figure 2A). Under standard conditions, the normal level of Hsp90 protein can be dramatically reduced without major impacts on growth rate, consistent with previous findings (Jiang et al, 2013; Picard et al, 1990).

**Figure 2.**
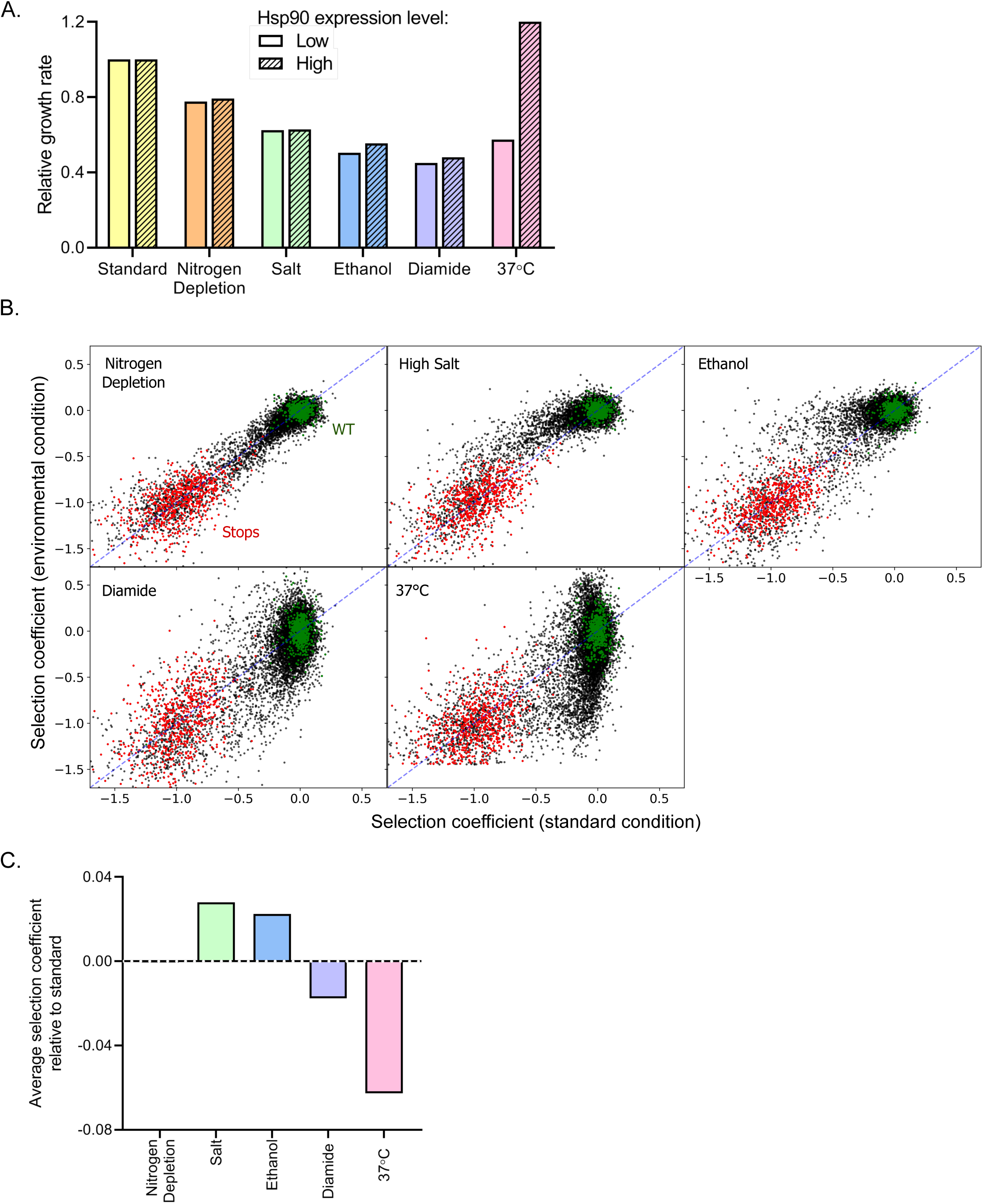
Impact of environmental stresses on yeast growth rates and selection on Hsp90 sequence. A. Growth rate of yeast with normal and reduced expression of Hsp90 protein in standard and stress conditions. B. Selection coeffcients of all Hsp90 amino acid variants in stress conditions compared to standard conditions. Wildtype amino acids are shown in green and stop codons are shown in red. Selection coeffcients were scaled to null (s=-1) for the average stop codon and neutral (s=0) for the average wildtype. C. The average selection coeffcient of all mutations relative to standard condtions, a metric of the strength of selection acting on Hsp90 sequence, in each stress condition.

We anticipated that Hsp90 would be required at increased levels for robust experimental growth in diamide, nitrogen starvation, ethanol, and high temperature (Gasch et al, 2000) based on the concept that cells increase expression level of genes in conditions where those gene products are needed at higher concentration. Consistent with this concept, reduced Hsp90 levels cause a marked decrease in growth rate at 37°C. However, Hsp90 protein levels had smaller impacts on growth rates under the other stress conditions, indicating that reliance on overall Hsp90 function does not increase dramatically in these conditions.

We quantified the growth rates of all Hsp90 single-mutant variants in each of the stress conditions as selection coefficients where 0 represents wildtype and −1 represents null alleles (Figure S2A-E, Table S1). We could clearly differentiate between the selection coefficients of wildtype synonyms and stop codons in all conditions (Figure 2B), and we normalized to these classes of mutations to facilitate comparisons between each condition. Of note, the observed selection coefficients of wildtype synonyms varied more in conditions of high temperature and diamide stress compared to standard (Figure S2F). We also note greater variation in the selection coefficients of barcodes for the same codon in the diamide and high temperature conditions (Figure S2G). We conclude that diamide and elevated temperature provided greater noise in our selection coefficient measurements. To take into account differences in signal to noise for each condition, we either averaged over large numbers of mutations or categorized selection coefficients as wildtype like, strongly deleterious, intermediate, or beneficial based on the distribution of wildtype synonyms and stop codons in each condition (see Materials and Methods and Figure S2H).

We compared selection coefficients of each Hsp90 variant in each stress condition to standard condition (Figure 2B&C). The stresses of 37°C and diamide tend to exaggerate the growth defects of many mutants compared to standard conditions, whereas high salt and ethanol tend to rescue growth defects (Figure 2B&C and S2I). According to the theory of metabolic flux (Dykhuizen et al, 1987; Kacser & Burns, 1981), gene products that are rate limiting for growth will be subject to the strongest selection. Accordingly, the relationship between Hsp90 function and growth rate should largely determine the strength of selection acting on Hsp90 sequence. Conditions where Hsp90 function is more directly linked to growth rate would be more sensitive to Hsp90 mutations than conditions where Hsp90 function can be reduced without changing growth rates (Bershtein et al, 2013; Jiang et al, 2013). The average selection coefficients are more deleterious in diamide and temperature stress compared to standard conditions. These findings are consistent with heat and diamide stresses causing a growth limiting increase in unfolded Hsp90 clients that is rate limiting for growth. In contrast, the average selection coefficients are less deleterious in ethanol and salt stress than in standard conditions, consistent with a decrease in the demand for Hsp90 function in these conditions. Due to the complex role Hsp90 plays in diverse signaling pathways in the cell, the different environmental stresses may differentially impact subsets of client proteins that cause distinct selection pressures on Hsp90 function.

### Structural analyses of environmental responsive positions

Altering environmental conditions had a pervasive influence on mutational effects along the sequence of Hsp90 (Figure 3A & S3A). We structurally mapped the average selection coefficient of each position in each condition relative to standard conditions as a measure of the sensitivity to mutation of each position under each environmental stress (Figure 3A). Many positions had mutational profiles that were responsive to a range of environments. Environmentally responsive positions with large changes in average selection coefficient in at least three conditions are highlighted on the Hsp90 structure in green in Figure 3B. Unlike the critical positions that cluster around the ATP binding site (Figure 1C), the environmentally responsive positions are located throughout all domains of Hsp90. Similar to critical residues, environmentally responsive positions are more conserved in nature compared to other positions in Hsp90 (Figure 3C), suggesting that the suite of experimental stress conditions tested captured aspects of natural selection pressures on Hsp90 sequence.

**Figure 3.**
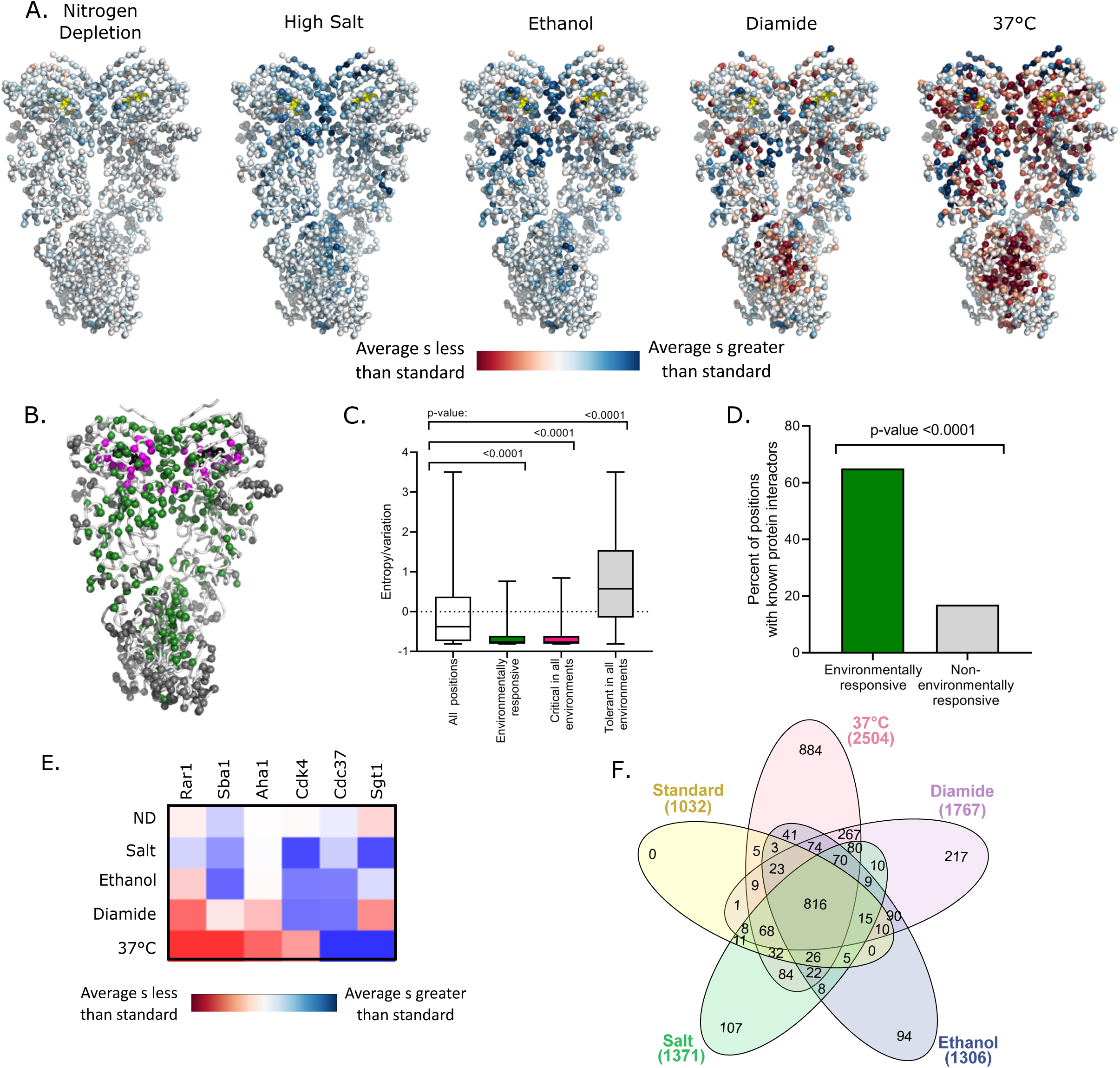
Environmental stresses place distinct selection pressures on Hsp90. A. The average selection coeffcient (s) at each position relative to standard conditions was mapped onto Hsp90 structure (Ali et al., 2006). B. Structural images indicating the location of positions that are critical for Hsp90 function in all conditions (magenta), positions that are environmentally responsive (green), and positions that are tolerant in all environments (gray). Critical residues have mean selection coeffcients that are null-like (within the distribution of stop codons) in all environments. Environmentally responsive positions have mean selection coeffcients that differed from standard in three or more environmental conditions by an amount greater than one standard deviation of wildtype synonyms. Tolerant residues are not shitied more than this cutoff in any environment. C. For different classes of positions, evolutionary variation was calculated as amino acid entropy at each position in Hsp90 sequences from diverse eukaryotes. Distributions are significantly different as measured by a two-sample Kolmogorov-Smirnov (KS) (All positions vs. environmentally responsive: N=678,55,p<0.0001,D-0.44; All positions vs. critical positions: N=678,55,p<0.0001,D=0.40; All positions vs. tolerant positions: N=678,136,p<0.0001,D=0.38) D. Fraction of different classes of mutations located at contact sites with binding partners. E. A heatmap of the average selection coeffcient for all positions at the stated interfaces relative to standard conditions in each environment. F. Venn diagram of deleterious mutations in different environmental conditions (Heberle, Meirelles, da Silva, Telles, & Minghim, 2015). Total number of deleterious mutants in each condition are stated in parentheses.

Hsp90 positions with environmentally responsive selection coefficients were enriched in binding contacts with clients, co-chaperones and intramolecular Hsp90 contacts involved in transient conformational changes (Figure 3D and S3B). About 65% of the environmentally responsive residues have been identified either structurally or genetically as interacting with binding partners (Ali et al, 2006; Bohen & Yamamoto, 1993; Genest et al, 2013; Hagn et al, 2011; Hawle et al, 2006; Kravats et al, 2018; Lorenz et al, 2014; Meyer et al, 2003; Meyer et al, 2004; Nathan & Lindquist, 1995; Retzlaff et al, 2009; Roe et al, 2004; Verba et al, 2016; Zhang et al, 2010), compared to about 15% of positions that were not responsive to stress conditions. While ATP binding and hydrolysis are the main structural determinants that constrain fitness in standard growth conditions, client and co-chaperone interactions have a larger impact on experimental fitness under stress conditions. Although the mean selection coefficients of mutations at the known client and co-chaperone binding sites are responsive to changes in environment, the direction of the shift of growth rate compared to standard conditions depends on the specific binding partner and environment (Figure 3E and S3C). This suggests that different environments place unique functional demands on Hsp90 that may be mediated by the relative affinities of different clients and co-chaperones. Consistent with these observations, we hypothesize that Hsp90 client priority is determined by relative binding affinity and that Hsp90 mutations can reprioritize clients that in turn impacts many signaling pathways.

### Constraint of mutational sensitivity at high temperature

We find that different environmental conditions impose unique constraints on Hsp90, with elevated temperature placing the greatest purifying selection pressure on Hsp90. Of the 2504 variants of Hsp90 that are deleterious when grown at 37°C, 884 of them (∼35%) are deleterious only in this condition (Figure 3F). We defined mutants that confer temperature sensitive (*ts*) growth phenotypes on cells as variants with selection coefficients within the distribution of wildtype synonyms in standard conditions and that of stop codons at 37°C. Based on this definition, 675 Hsp90 amino acid changes (roughly 5% of possible changes) were found to be temperature sensitive (Figure 4A). We sought to understand the physical underpinnings of this large set of Hsp90 *ts* mutations.

**Figure 4.**
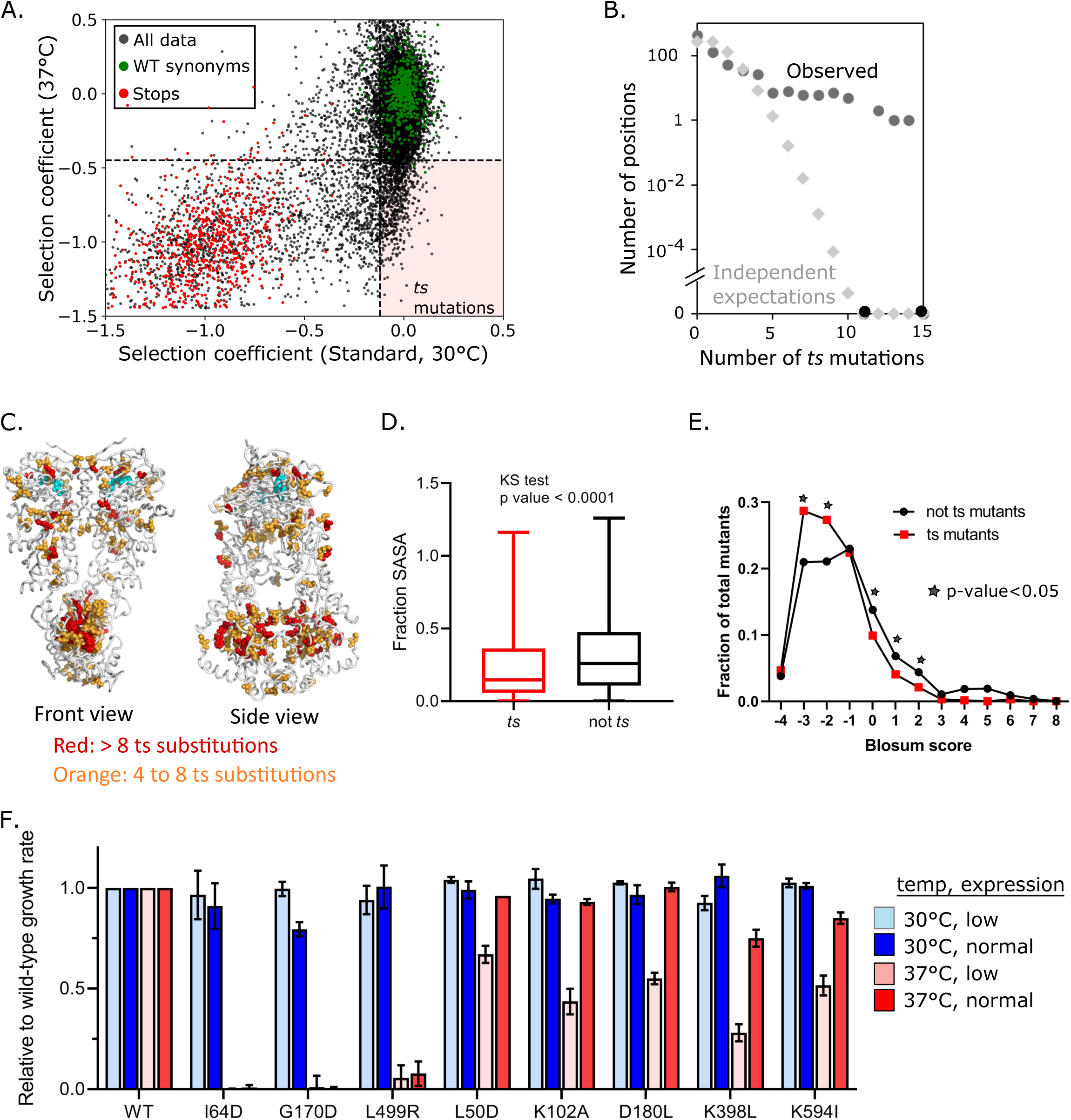
Abundance and mechanism of temperature sensitive mutations in Hsp90. A. Temperature sensitive (*ts*) amino acid variants were identified that supported wildtype-like growth at standard (30°C) temperature, but were null-like at 37°C in bulk competitions. B. Observed *ts* varaints tend to cluster at specific positions in Hsp90 in a significant manner compared to what would be expected if they occurred independently. C. Mapping positions with multiple *ts* variants onto Hsp90 structure. D. Solvent accessible surface area (SASA) of positions with *ts* mutations compared to positions lacking *ts* mutations. A two sample KS test showed significant differences in distributions (N=663, 11762, p<0.0001, D=0.1735) E. Amino acid similarity to the wildtype was estimated as the Blosum score for *ts* and non-*ts* variants. A two-proportion z-test was performed on each pair for each Blosum score and their p-values were adjusted using Benjamini-Hochberg adaptive step-up procedure. F. Growth rate of a panel of individual Hsp90 *ts* variants analyzed in isolation.

We examined Hsp90 *ts* mutations for structural and physical patterns. We found that *ts* mutations tended to concentrate at hotspots (Figure 4B). These hotspots were spread across all three domains of Hsp90 (Figure 4C). The largest cluster of hotspots occurred in the C domain of Hsp90. The C domain forms a constitutive homodimer that is critical for function (Wayne & Bolon, 2007). Of note, homo-oligomerization domains may have a larger *ts* potential because all subunits contribute to folding and dimerization essentially multiplying the impacts of mutations (Lynch, 2013). To explore the physical underpinnings of *ts* mutations we examined if they were buried in the structure or surface exposed. Mutations at buried residues tend to have a larger impact on protein folding energy compared to surface residues (Chakravarty & Varadarajan, 1999). Consistent with the idea that many *ts* mutations may disrupt protein folding at elevated temperature, substitutions that confer a *ts* phenotype are enriched in buried residues (Figure 4D). Also consistent with this idea, *ts* mutations tend to have negative Blosum scores (Figure 4E), a hallmark of disruptive amino acid changes.

Because growth at elevated temperatures requires higher levels of Hsp90 protein (Borkovich et al, 1989), some *ts* mutations are likely due to a reduced function that is enough for growth at standard temperature, but is insufficient at 37°C (Nathan & Lindquist, 1995). We reasoned that we could distinguish these mutants by examining how growth rate depended on the expression levels of Hsp90. We expect that destabilizing mutants that cause Hsp90 to unfold at elevated temperature would not support efficient growth at 37°C independent of expression levels. In contrast, we expect mutants that reduce Hsp90 function to exhibit an expression-dependent growth defect at 37°C. We tested a panel of *ts* mutations identified in the bulk competitions at high and low expression levels (Figure 4F). The dependence of growth rate at 37°C on expression level varied for different Hsp90 *ts* variants. The I64D, G170D and L499R Hsp90 mutants have no activity at 37°C irrespective of expression levels. These disruptive substitutions at buried positions likely destabilize the structure of Hsp90. In contrast, increasing the Hsp90 expression levels at least partially rescued the growth defect for five *ts* variants (L50D, K102A, D180L, K398L, K594I), indicating that these variants do not providing enough Hsp90 function for robust growth at elevated temperature. All five of these expression dependent *ts* variants were located at surface positions, indicating that the location of *ts* mutations can delineate different mechanistic classes.

### Hsp90 potential for adaptation to environmental stress

Numerous Hsp90 variants provided a growth benefit compared to the wildtype sequence in stress conditions. The largest number of beneficial variants in Hsp90 occurred in high temperature and diamide conditions (Figure 5A). Multiple lines of evidence indicate that these mutants are truly beneficial variants and not simply measurement noise. First, the beneficial amino acids generally exhibited consistent selection coefficients among synonymous variants (Figure S5A). Second, adaptive mutants in diamide and high temperature cluster at certain positions in a significant manner (see below). Finally, we confirmed the increased growth rate at elevated temperature of a panel of variants analyzed in isolation (Figure S5B). Beneficial mutations in elevated temperature and diamide often clustered at specific positions in Hsp90 (Figure 5B), indicating that the wildtype amino acids at these positions are far from optimum for growth in these conditions. In contrast, the apparent beneficial mutations in other conditions did not tend to cluster at specific positions (Figure S5C).

**Figure 5.**
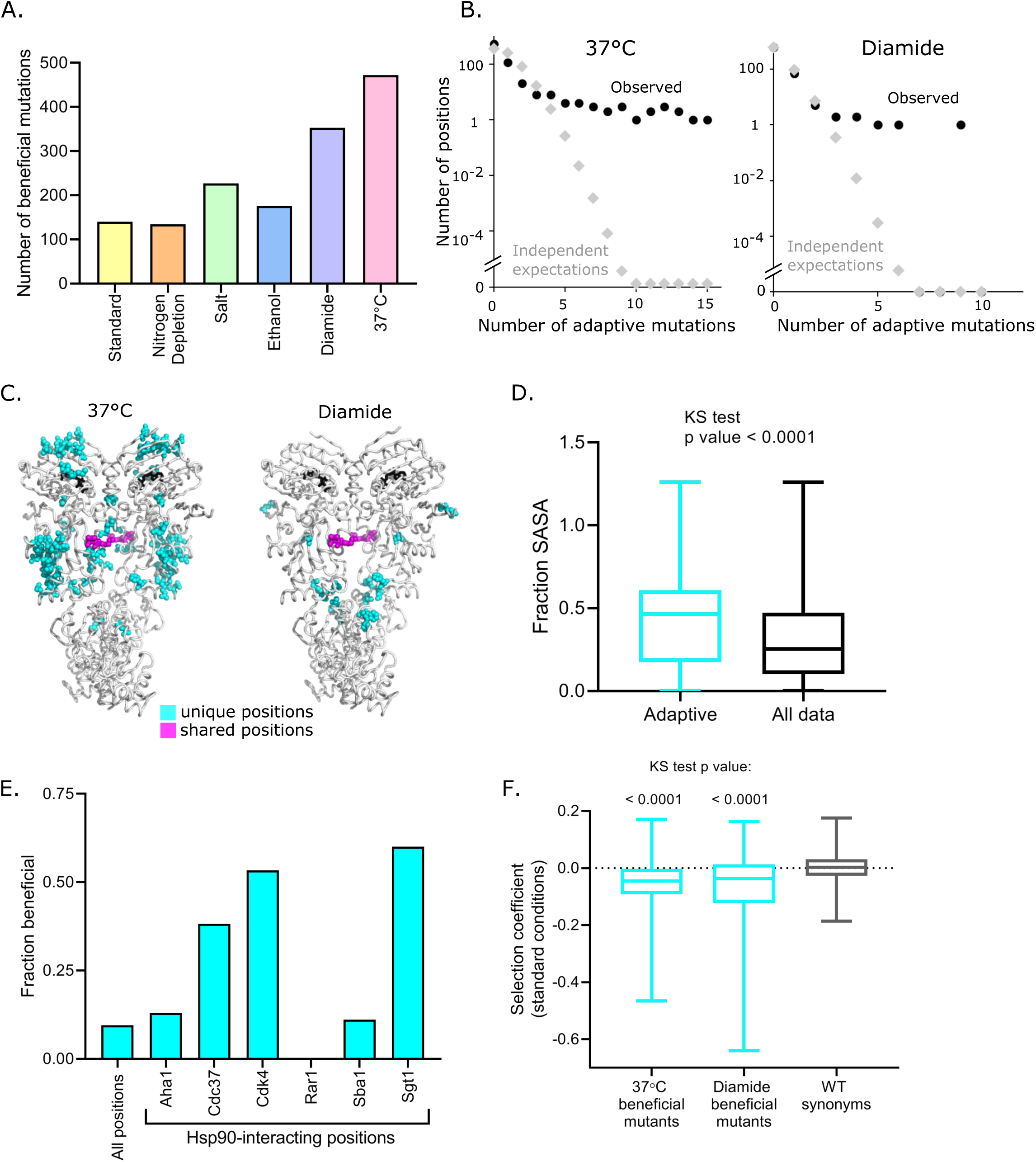
Beneficial variants in diamide and elevated temperature conditions. A. Number of beneficial mutations identified in each condition based on selection coeffcients more than two standard deviations greater than wildtype synonyms. B. Distribution of the number of beneficial mutations at the same position in both 37°C (leti) and diamide (right) conditions. C. Location of positions with four or more beneficial mutations. Positions that are unique to diamide or 37°C are shown in cyan and two shared positions are shown in magenta. D. The solvent accessible surface area of beneficial positions at 37°C and in diamide compared to all positions. Distributions are significantly different as measured by a two-sample KS test (N=463,12393,p<0.0001,D=0.3809) E. The fraction of Hsp90 positions at interfaces that were beneficial in 37°C and diamide conditions. F. Selection coeffcients in standard conditions for beneficial mutations at 37 °C and in diamide compared to wildtype synonyms. (KS test; 37°C vs. WT synonyms: N=463,660,p<0.0001,D=0.3281; Diamide vs. WT synonyms: N=276,660,p<0.0001,D=0.3809)

To obtain a more general picture of the potential for adaptation derived from the full fitness distributions, we used Fisher’s Geometric model (FGM) (Fisher, 1930). According to FGM, populations evolve in an *n*-dimensional phenotypic space, through random single step mutations, and any such mutation that brings the population closer to the optimum is considered beneficial. An intuitive hypothesis derived from FGM is that the potential for adaptation in a given environment (that is the availability of beneficial mutations) depends on the distance to the optimum. In order to estimate the distance to the optimum *d*, we adopted the approach by Martin and Lenormand and fitted a displaced gamma distribution to the neutral and beneficial mutations for each environment (Martin & Lenormand, 2006). We observed that the yeast populations were furthest from the optimum in elevated temperature and diamide (*d*=0.072 and 0.05, respectively), followed by nitrogen deprivation (*d*=0.023), high salinity and ethanol (*d*=0.021) and standard (d=0.014). This suggests that exposure to elevated temperature and diamide results in the largest potential for adaptation and is consistent with the observation of the largest proportions of beneficial mutations in these environments. Interestingly, previous results from a 9-amino-acid region in Hsp90 indicated that there was very little potential for adaptation at high temperature (36°C) as compared with high salinity (Hietpas, 2013). This apparent contradiction between results from the full Hsp90 sequence and the 582-590 region indicates that a specific region of the protein may be already close to its functional optimum in a specific environment, whereas there is ample opportunity for adaptation when the whole protein sequence is considered.

In diamide and elevated temperature, the clustered beneficial positions were almost entirely located in the ATP-binding domain and the middle domain (Figure 5C), both of which make extensive contacts with clients and co-chaperones (Ali et al, 2006; Meyer et al, 2003; Meyer et al, 2004; Roe et al, 2004; Verba et al, 2016; Zhang et al, 2010). Beneficial mutations in elevated temperature and diamide conditions were preferentially located on the surface of Hsp90 (Figure 5D) at positions accessible to binding partners. Analyses of available Hsp90 complexes indicate that beneficial positions were disproportionately located at known interfaces with co-chaperones and clients (Figure 5E). Clustered beneficial mutations are consistent with disruptive mechanisms because a number of different amino acid changes can lead to disruptions, whereas a gain of function is usually mediated by specific amino acid changes. Amino acids that are beneficial in diamide and elevated temperature tend to exhibit deleterious effects in standard conditions (Figure 5F), consistent with a cost of adaptation. We conjecture that the clustered beneficial mutations are at positions that mediate the binding affinity of subsets of clients and co-chaperones and that disruptive mutations at these positions can lead to re-prioritization of multiple clients. The priority or efficiency of Hsp90 for sets of clients can in turn impact most aspects of physiology because Hsp90 clients include hundreds of kinases that influence virtually every aspect of cell biology.

In the first ten amino acids of Hsp90 we noted a large variation in the selection coefficients of synonymous mutations at elevated temperature (Figure S5D). These synonymous mutations were only strongly beneficial at high temperature where Hsp90 protein levels are limiting for growth. Analysis of an individual clones confirms that synonymous mutations at the beginning of Hsp90 that were beneficial at high temperature were expressed at higher level in our plasmid system (Figure S5E, S5F). These results are consistent with a large body of research showing that mRNA structure near the beginning of coding regions often impacts translation efficiency (Li, 2015; Plotkin & Kudla, 2011; Tuller et al, 2010), and that adaptations can be mediated by changes in expression levels (Lang & Desai, 2014). Outside of the first ten amino acids, we did not observe large variation in selection coefficients of synonymous mutations.

### Natural selection favors Hsp90 variants that are robust to environment

We next examined how experimental protein fitness maps compared with the diversity of Hsp90 sequences in current eukaryotes. We analyzed Hsp90 diversity in a set of 267 sequences from organisms that broadly span across eukaryotes. We identified 1750 amino acid differences in total that were located at 499 positions in Hsp90. We examined the experimental growth effects of the subset of amino acids that were observed in nature. While the overall distribution of selection coefficients in all conditions was bimodal with peaks around neutral (s=0) and null (s=-1), the natural amino acids were unimodal with a peak centered near neutral (Figure 6A). The vast majority of natural amino acids had wildtype-like fitness in all conditions studied here (Figure 6B&C). Whereas naturally occurring amino acids in Hsp90 were rarely deleterious in any experimental condition, they were similarly likely to provide a growth benefit compared to all possible amino acids (5%). This observation indicates that condition-dependent fitness benefits are not a major determinant of natural variation in Hsp90 sequences. Instead, our results indicate that natural selection has favored Hsp90 substitutions that are robust to multiple stressful conditions (Figure 6D).

**Figure 6.**
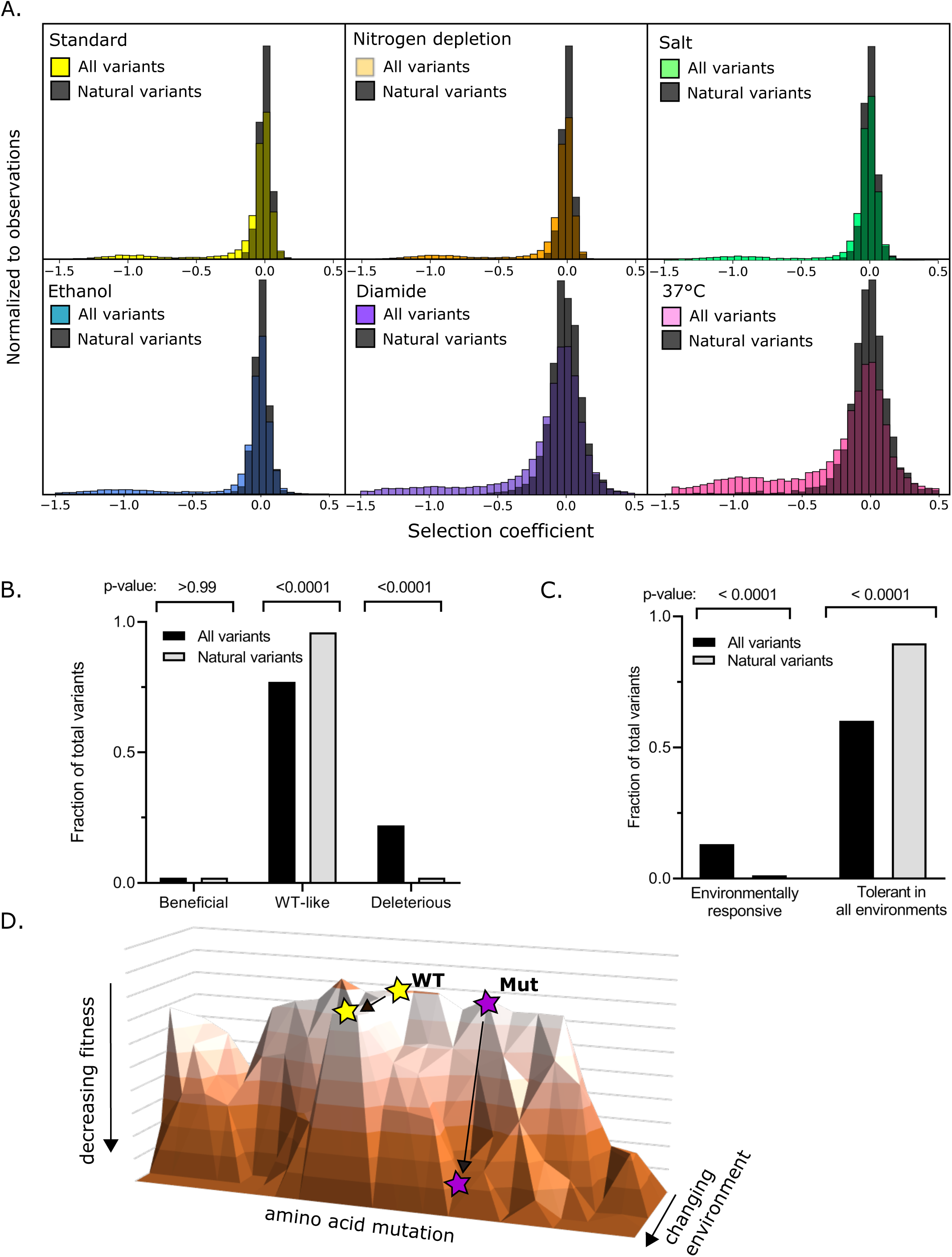
Experimental growth effects of natural amino acid variants of Hsp90. A. The distribution of selection coeffcients of natural variants compared to all variants in each environmental condition. B. Across all environments, the fraction of natural variants compared to all variants that were beneficial, wildtype like, or deleterious. C. The fraction of natural variants compared to all variants that were environmentally responsive or tolerant in all environments. Categories were defined as in Figure 3B. D. Landscape model indicating that natural variants of Hsp90 tend to support robust growth under a variety of stress conditions.

Epistasis may provide a compelling explanation for the naturally occurring amino acids that we observed with deleterious selection coefficients. Analyses of Hsp90 mutations in the context of likely ancestral states has demonstrated a few instances of historical substitutions with fitness effects that depend strongly on the Hsp90 sequence background (Starr et al, 2018). Indeed, many of the natural amino acids previously identified with strong epistasis (E7A, V23F, T13N) are in the small set of natural amino acids with deleterious effects in at least one condition. Further analyses of natural variants under diverse environmental conditions will likely provide insights into historical epistasis and will be the focus of future research.

## DISCUSSION

In this study, we analyzed the protein-wide distribution of fitness effects of Hsp90 across standard and five stress conditions. We found that environment has a profound effect on the fates of Hsp90 mutations. Each environmental stress varies in the strength of selection on Hsp90 mutations; heat and diamide increase the strength of selection and ethanol and salt decrease the strength of selection. While proximity to ATP is the dominant functional constraint in standard conditions, the influence of client and co-chaperone interactions on growth rate dramatically increases under stress conditions. Additionally, beneficial mutations cluster at positions that mediate binding to clients and cochaperones. The fact that different Hsp90 binding partners have distinct environmental dependencies suggests that Hsp90 can reprioritize clients that in turn impacts many downstream signaling pathways.

Our results demonstrate that mutations to Hsp90 can have environment-dependent effects that are similar to the stress induced changes to the function of wildtype Hsp90 that have been shown to contribute to new phenotypes (Jarosz et al, 2010). The low frequency of environment-dependent amino acids in Hsp90 from extant eukaryotes indicates that this type of evolutionary mechanism is rare relative to drift and other mechanisms shaping Hsp90 sequence diversity.

We observed distinct structural trends for mutations that provide environment-dependent costs and benefits. Many mutations in Hsp90 caused growth defects at elevated temperature where Hsp90 function is limiting for growth. These temperature sensitive mutations tended to be buried and in the homodimerization domain, consistent with an increased requirement for folding stability at elevated temperatures. In contrast, beneficial mutations tended to be on the surface of Hsp90 and at contact sites with binding partners, suggesting that change-of-function mutations may be predominantly governed by alterations to binding interactions.

Importantly, our results demonstrate that while mutations to Hsp90 can provide a growth advantage in specific environmental conditions, naturally occurring amino acids in Hsp90 tend to support robust growth over multiple stress conditions. The finding of beneficial mutations in Hsp90 in specific conditions suggests that similar long-term stresses in nature can lead to positive selection on Hsp90. Consistent with previous work (Hietpas, 2013), we found that experimentally beneficial mutations tended to have a fitness cost in alternate conditions (Figure 5F). This indicates that natural environments which fluctuate among different stresses would reduce or eliminate positive selection on Hsp90. Therefore, our results suggest that natural selection on Hsp90 sequence has predominantly been governed by strong purifying selection integrated over multiple stressful conditions. Taken together, these results support the hypothesis that natural populations might experience a so-called “micro-evolutionary fitness seascape” (Mustonen & Lassig, 2009), in which rapidly fluctuating environments result in a distribution of quasi-neutral substitutions over evolutionary time scales.

## ACKNOWLEDGEMENTS

Thanks to Tyler Starr for providing the alignment of Hsp90 sequences used to assess natural variation. This work was supported by grants from the National Institutes of Health (R01-GM112844 to D.N.A.B. and F32-GM119205 to J.M.F). I.F. was supported by a postdoctoral fellowship from the FCT (Fundação para a Ciência e a Tecnologia) within the project JPIAMR/0001/2016. C.B. is grateful for support from EMBO Installation Grant IG4152 and ERC Starting Grant 804569 - FIT2GO.

## MATERIALS AND METHODS

### Generating Mutant Libraries

A library of Hsp90 genes was saturated with single point mutations using oligos containing NNN codons as previously described (Hietpas et al, 2012). The resulting library was pooled into 12 separate 60 amino acid long sub-libraries (amino acids 1-60, 61-120 etc.) and combined via Gibson Assembly (NEB) with a linearized p414ADHΔter Hsp90 destination vector. To simplify sequencing steps during bulk competition, each variant of the library was tagged with a unique barcode. For each 60 amino acid sub-library, a pool of DNA constructs containing a randomized 18-bp barcode sequence (N18) was cloned 200 nt downstream from the Hsp90 stop codon via restriction digestion, ligation, and transformation into chemically competent E. coli with the goal of each mutant being represented by 10-20 unique barcodes.

### Barcode Association of Library Variants

We added barcodes and associated them with Hsp90 variants essentially as previously described (Starr et al, 2018). To associate barcodes with Hsp90 variants, we performed paired-end sequencing of each 60 amino acid sub-library using a primer that reads the N18 barcode in one read and a primer unique to each sub-library that anneals upstream of the region containing mutations. To facilitate efficient Illumina sequencing, we generated PCR products that were less than 1kb in length for sequencing. We created shorter PCR products by generating plasmids with regions removed between the randomized regions and the barcode. To remove regions from the plasmids, we performed restriction digest with two unique enzymes, followed by blunt ending with T4 DNA polymerase (NEB) and plasmid ligation at a low concentration (3 ng/μL) to favor circularization over bimolecular ligations. The resulting DNA was re-linearized by restriction digest, and amplified with 11 cycles of PCR to generate products for Illumina sequencing. The resulting PCR products were sequenced using an Illumina MiSeq instrument with asymmetric reads of 50 bases for Read1 (barcode) and 250 bases for Read2 (Hsp90 sequence). After filtering low-quality reads (Phred scores <10), the data was organized by barcode sequence. For each barcode that was read more than three times, we generated a consensus of the Hsp90 sequence that we compared to wildtype to call mutations.

### Bulk Growth Competitions

Equal molar quantities of each sub-library were mixed to form a pool of DNA containing the entire Hsp90 library with each codon variant present at similar concentration. The plasmid library was transformed using the lithium acetate procedure into the DBY288 Hsp90 shutoff strain essentially as previously described (Jiang et al, 2013). Sufficient transformation reactions were performed to attain ∼5 million independent yeast transformants representing a 5-fold sampling for the average barcode and 50 to 100-fold sampling for the average codon variant. Following 12 hours of recovery in SRGal (synthetic 1% raffinose and 1% galactose) media, transformed cells were washed five times in SRGal-W (SRGal lacking tryptophan) media to remove extracellular DNA, and grown in SRGal-W media at 30°C for 48 h with repeated dilution to maintain the cells in log phase of growth. This yeast library were was supplemented with 20% glycerol, aliquoted and slowly frozen in a −80°C freezer.

For each competition experiment, an aliquot of the frozen yeast library cells was thawed at 37°C. Viability of the cells was accessed before and after freezing and was determined to be greater than 90% with this slow freeze, quick thaw procedure. Thawed cells were amplified in SRGal-W for 24 hours, and then shifted to shutoff conditions by centrifugation, washing, and resuspension in 300 mL of synthetic dextrose lacking tryptophan (SD-W) for 12 hours at 30°C. At this point, cells were split and transferred to different conditions including: Standard (SD-W, 30°C), Nitrogen depletion (SD-W with limiting amounts of ammonium sulfate, 0.0125%, 30°C), High salt (SD-W with 0.8 M NaCl, 30°C), Ethanol (SD-W with 7.5% ethanol, 30°C, Diamide (SD-W with 0.85 mM diamide, 30°C), or high temperature (SD-W, 37°C). We collected samples of ∼10^8^ cells at eight time points over a period of 36 hours and stored them at −80°C. Cultures were maintained in log phase by regular dilution with fresh media, maintaining a population size of 10^9^ or greater throughout the bulk competition. Bulk competition from the standard condition were conducted in technical duplicates from the frozen yeast library.

### DNA Preparation and Sequencing

We isolated plasmid DNA from each bulk competition time point as described (Jiang et al, 2013). Purified plasmid was linearized with AscI. Barcodes were amplified by 19 cycles of PCR using Phusion polymerase (NEB) and primers that add Illumina adapter sequences and an 8 bp identifier sequence used to distinguish libraries and time points. The identifier sequence was located at positions 91-98 relative to the illumine primer and the barcode was located at positions 1-18. PCR products were purified two times over silica columns (Zymo Research) and quantified using the KAPA SYBR FAST qPCR Master Mix (Kapa Biosystems) on a Bio-Rad CFX machine. Samples were pooled and sequenced on an Illumina NextSeq instrument in single-end 100 bp mode.

### Analysis of Bulk Competition Sequencing Data

Illumina sequence reads were filtered for Phred scores >20 and strict matching of the sequence to the expected template and identifier sequence. Reads that passed these filters were parsed based on the identifier sequence. For each condition/time-point identifier, each unique N18 read was counted. The unique N18 count file was then used to identify the frequency of each mutant using the variant-barcode association table. To generate a cumulative count for each codon and amino acid variant in the library, the counts of each associated barcode were summed. To reduce experimental noise, selection coefficients were not calculated for variants with less than 100 reads at the 0 time point (Boucher et al, 2014). The average variant at the 0 time point had approximately 500 reads.

### Determination of Selection Coefficient

Selection coefficients were estimated using empiricIST (Fragata et al, 2018), a software package developed based on a previously published Markov Chain Monte Carlo (MCMC) approach (Bank et al, 2014). Briefly, we estimated individual growth rates and initial population sizes relative to the wildtype sequence simultaneously, based on a model of exponential growth and multinomial sampling of sequencing reads independently at each time point. For each mutant we obtained 10,000 posterior samples for the growth rate and initial population using a Metropolis-Hastings algorithm. The resulting growth rate estimates correspond to the median of 1,000 samples of the posterior. Subsequently, selection coefficients (s) were scaled so that the average stop codon in each environmental condition represented a null allele (s=-1). For the second replicate in standard conditions, we noted a small fitness defect (s≈-0.05) for wildtype synonyms at positions 679-709 relative to other positions. We do not understand the source of this behavior, and chose to normalize to wildtype synonyms from 1-678 for this condition and to exclude positions 679-709 from analyses that include the second replicate of standard conditions. We did not observe this behavior in any other condition. Variants were categorized as having wildtype-like, beneficial, intermediate, or deleterious fitness based on the comparison of their selection coefficients with the distribution of wild-type synonyms and stop codons in each condition (Figure S2H) in the following manner; Wildtype-like: variants with selection coefficients within two standard deviations (SD) of the mean of wildtype synonyms; Beneficial: variants with selection coefficients above 2 SD of wildtype synonyms; Strongly deleterious: variants with selection coefficients within 2 SD of stop codons; Intermediate: variants with selection coefficients between those of stop-like and wildtype-like.

### Yeast growth analysis

Individual Hsp90 variants were generated and analyzed essentially as previously described (Jiang et al, 2013). Variants were generated by site directed mutagenesis and transformed into DBY288 cells. Selected transformed colonies were grown in liquid SRGal-W media to mid-log phase at 30°C, washed three times and grown in shutoff media (SD-W) at either 30C or 37C. After sufficient time to stall the growth of control cells lacking a rescue copy of Hsp90 (∼16 hours), cell density was monitored based on absorbance at 600 nm over time and fit to an exponential growth curve to quantify growth rate.

### Natural variation in Hsp90 sequence

We analyzed sequence variation in a previously described alignment of Hsp90 protein sequences from 261 eukaryotic species that broadly span a billion years of evolutionary distance (Starr et al, 2018).

---

**Supplementary Figure 1.**
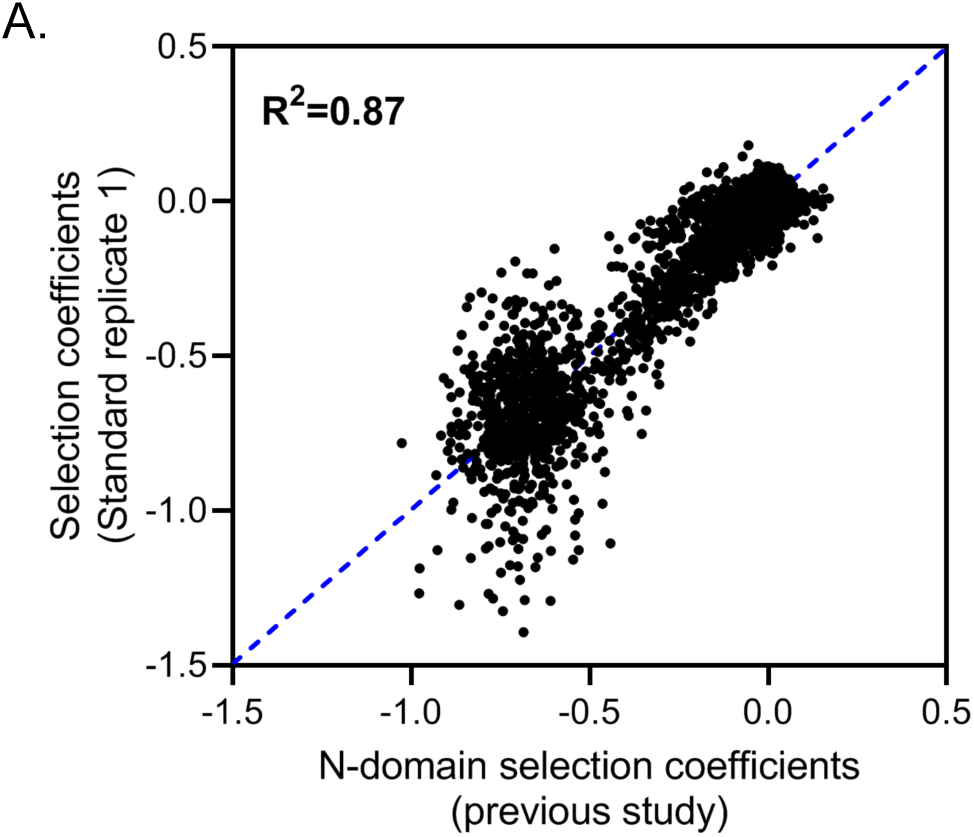
A. Measurement of selection coeffcients for positions 2-220 in this study correlated strongly (R^2^=0.87) with estimates of the Hsp90 N-domain in a previous study (Mishra, Flynn & Bolon, 2016), indicating that biological replicates show high reproducibility.

**Supplementary Figure 1.**
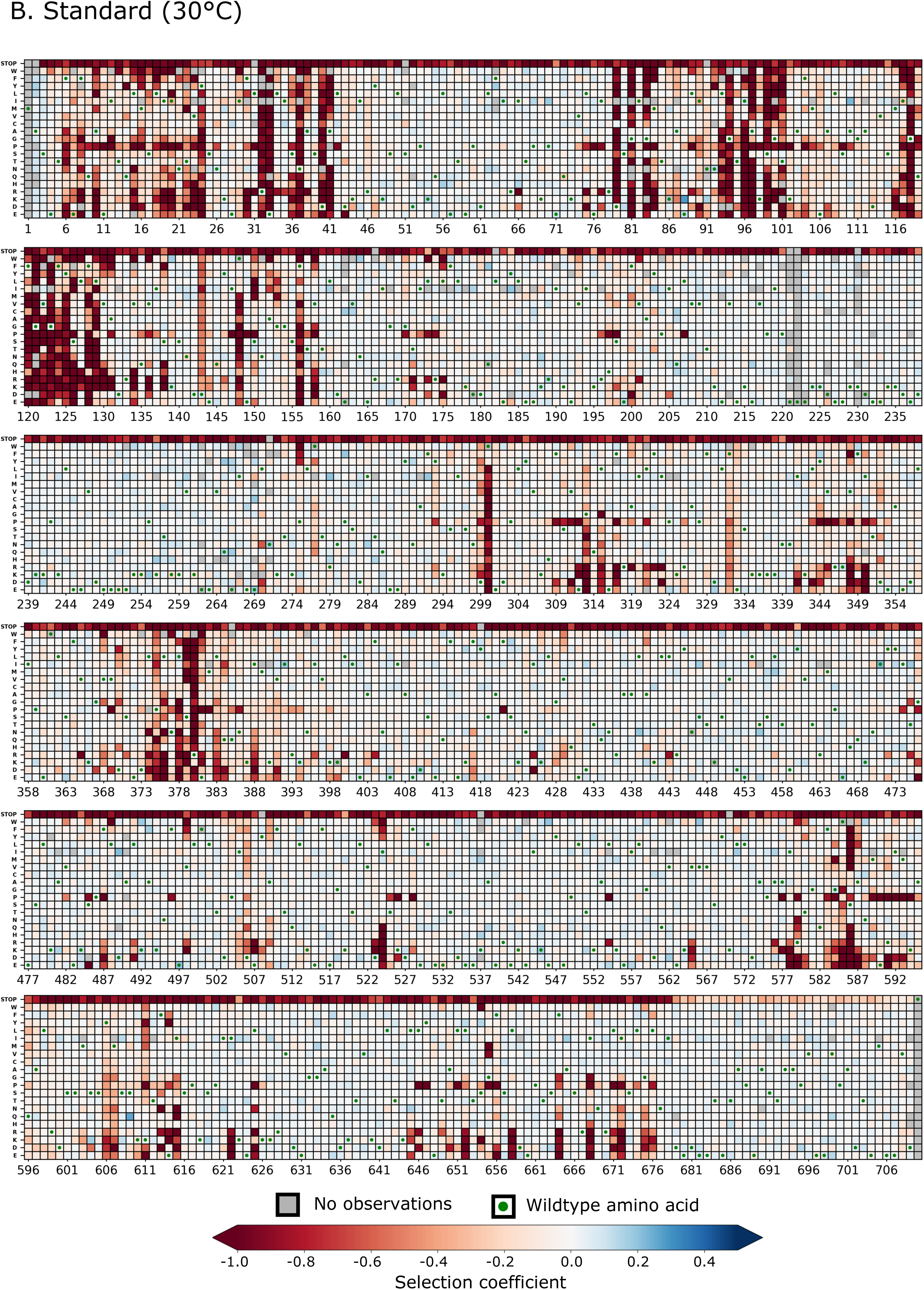
B. Heatmap representation of the fitness map observed for single amino acid changes across amino acids 2-709 of Hsp90 in standard (30°C) conditions.

**Supplementary Figure 1.**
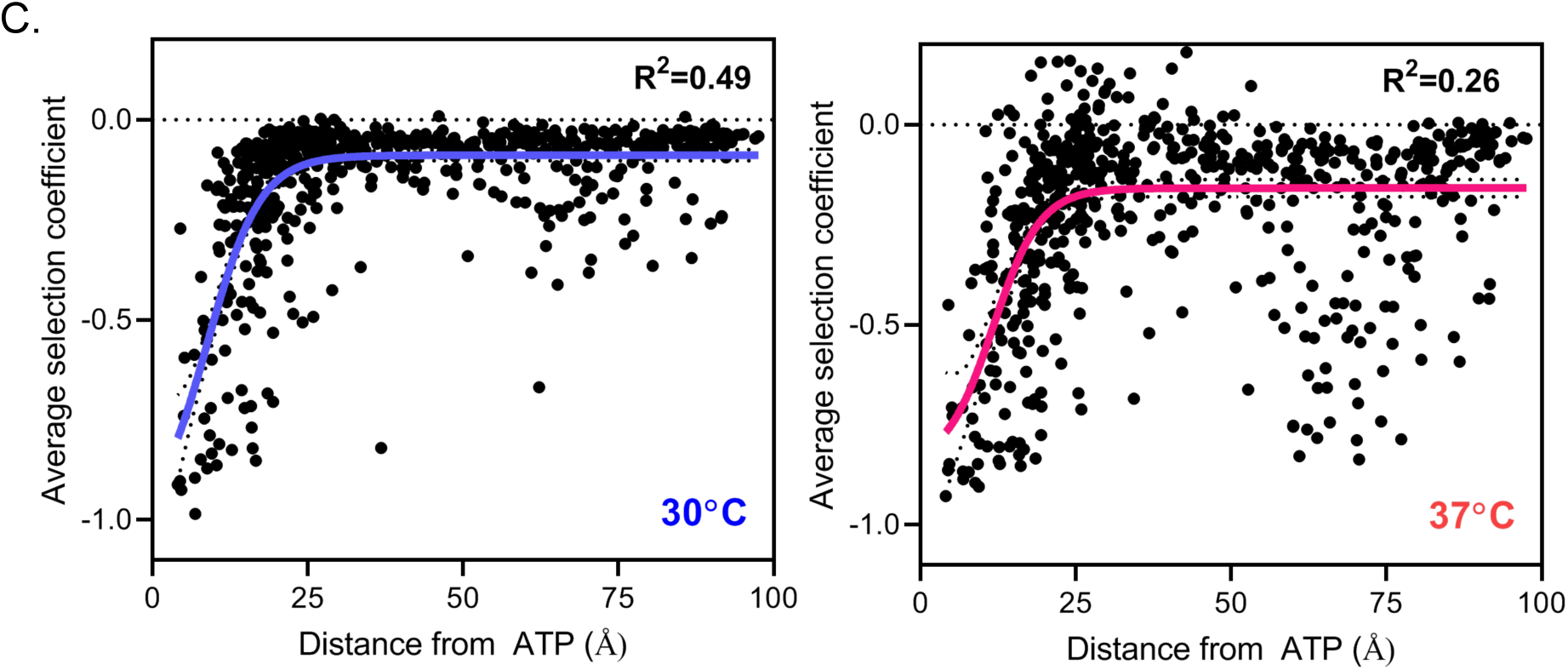
C. Average selection coeffcient at each position (a measure of mutational sensitivity) correlates with distance to ATP to a much higher degree in the 30°C data set (leti) compared to the 37°C data set (right).

**Supplementary Figure 2.**
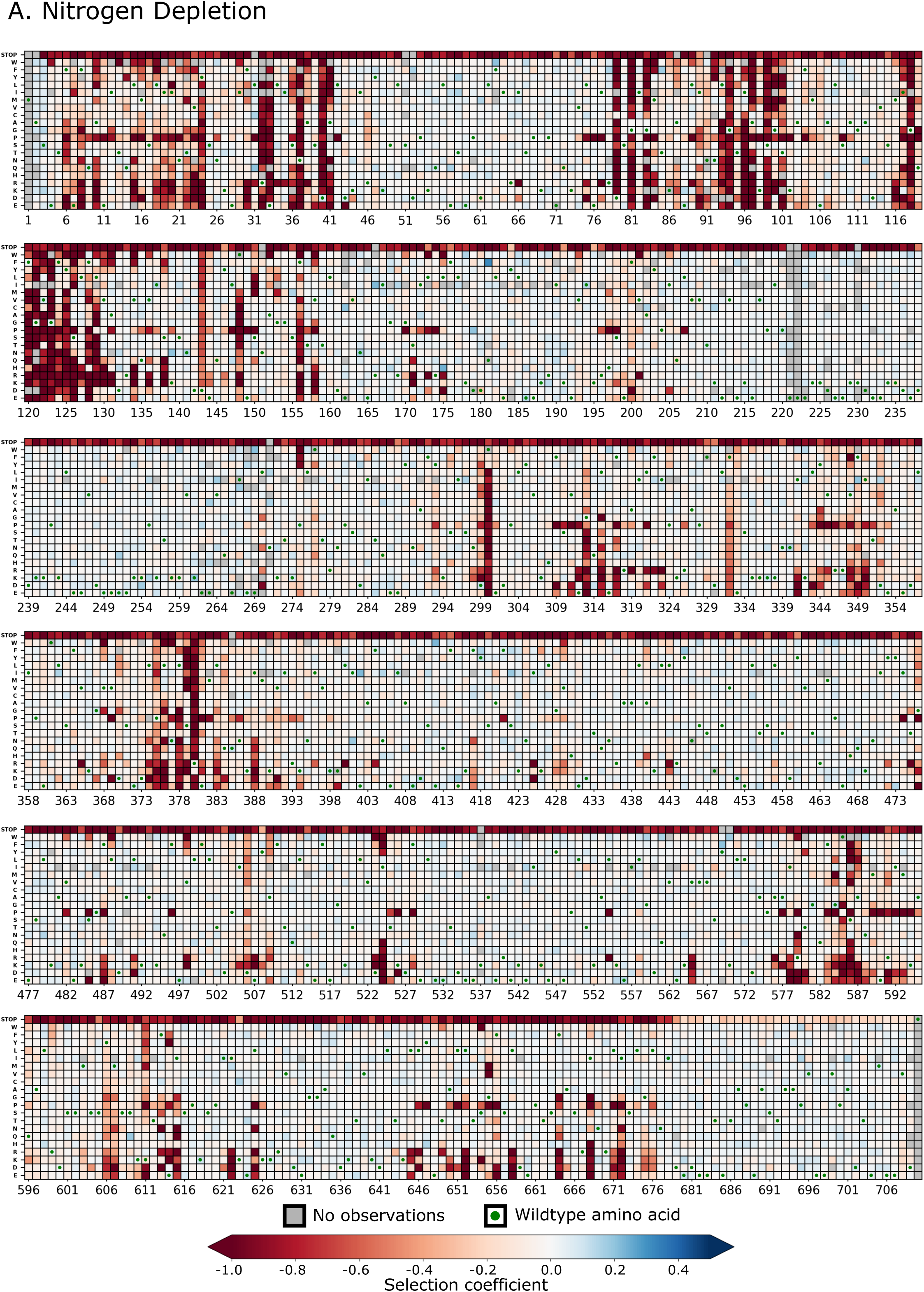
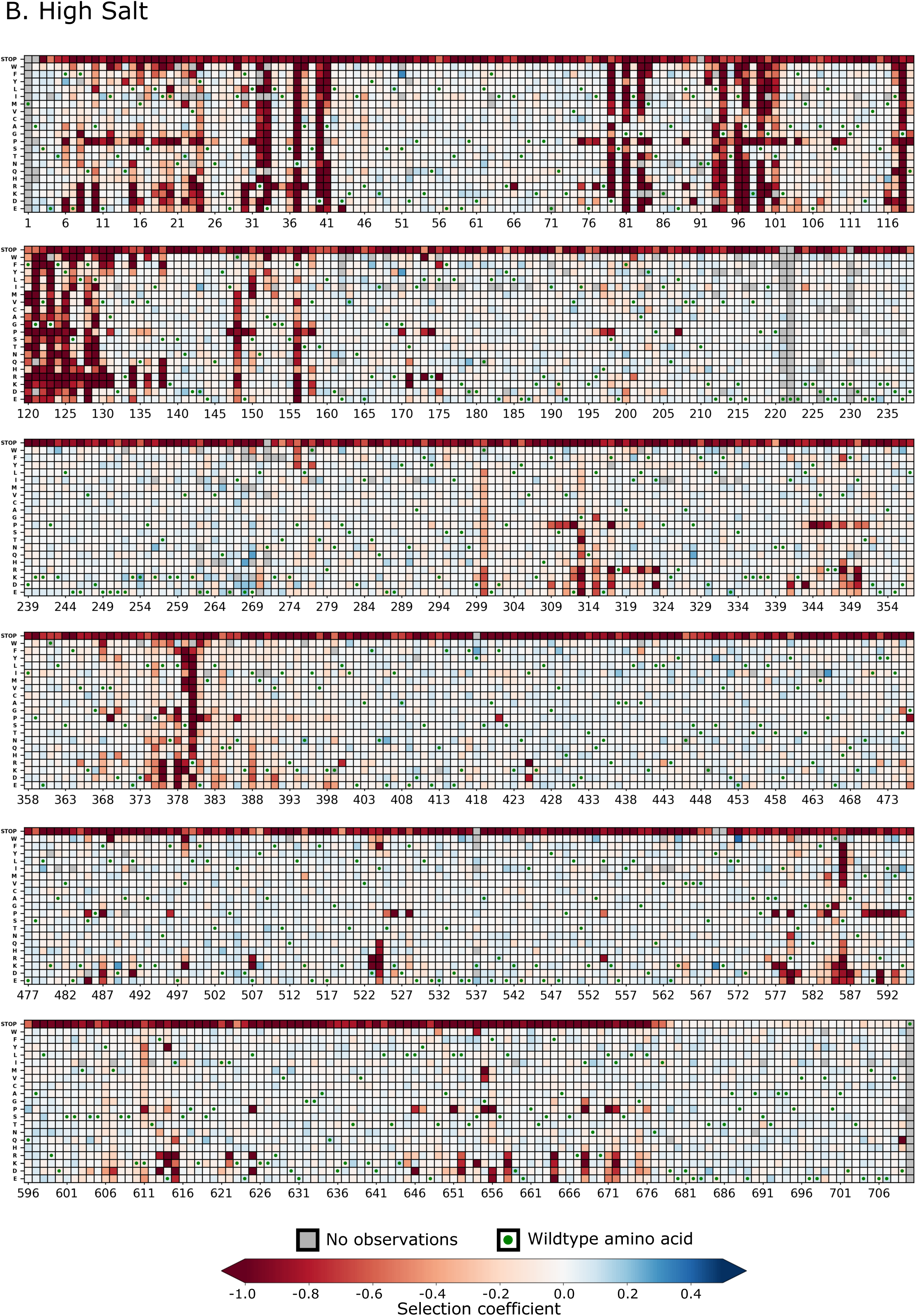
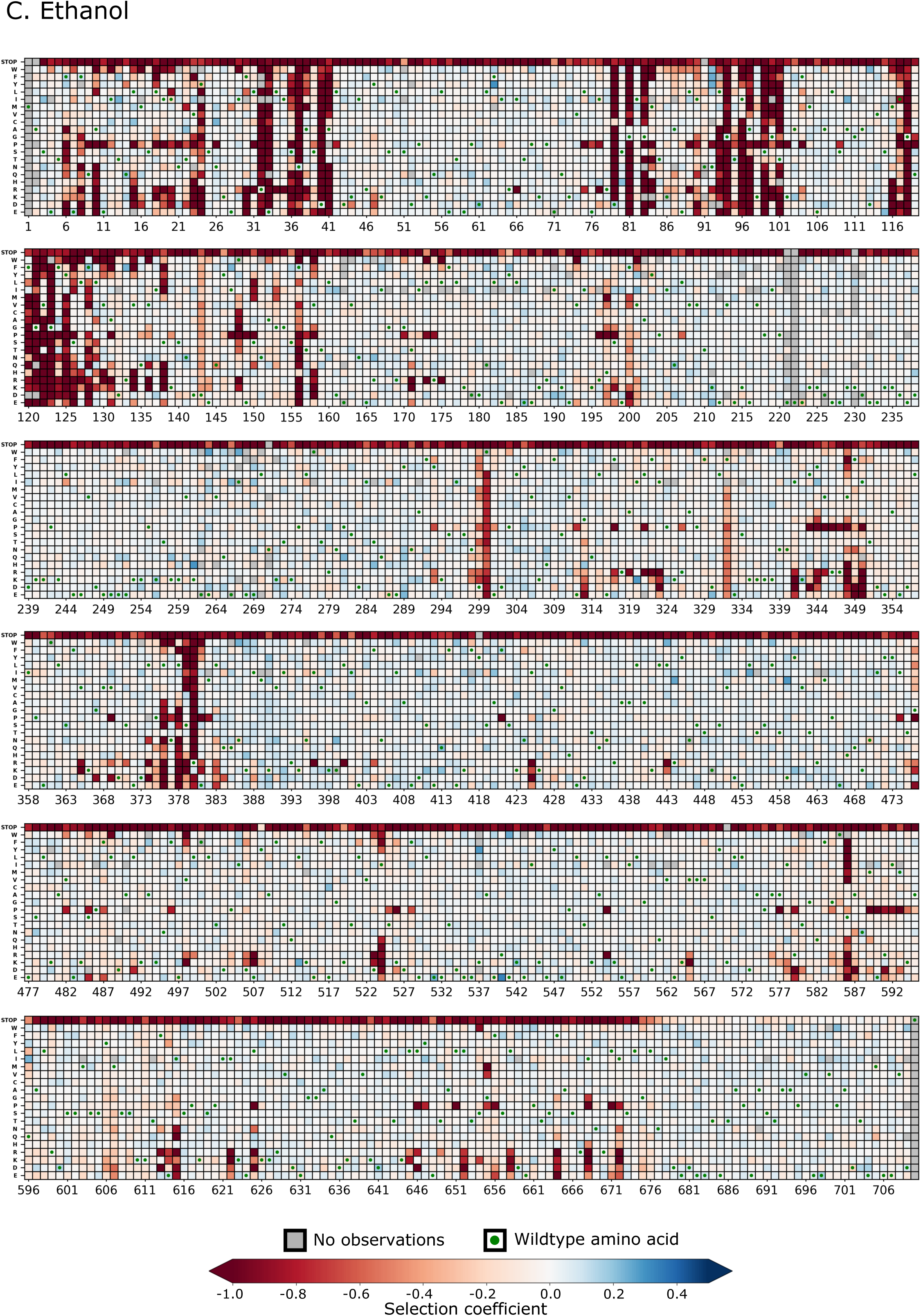
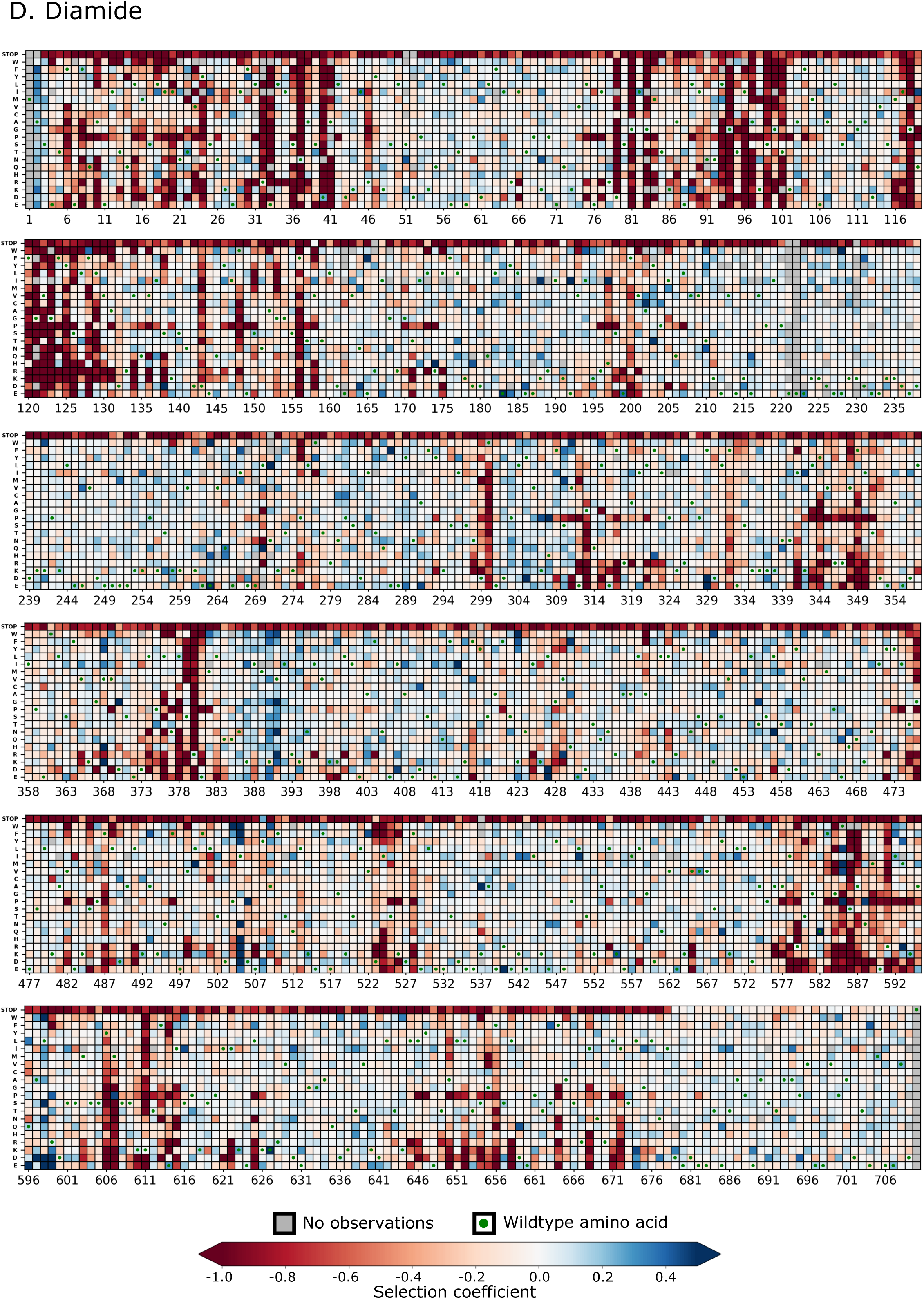
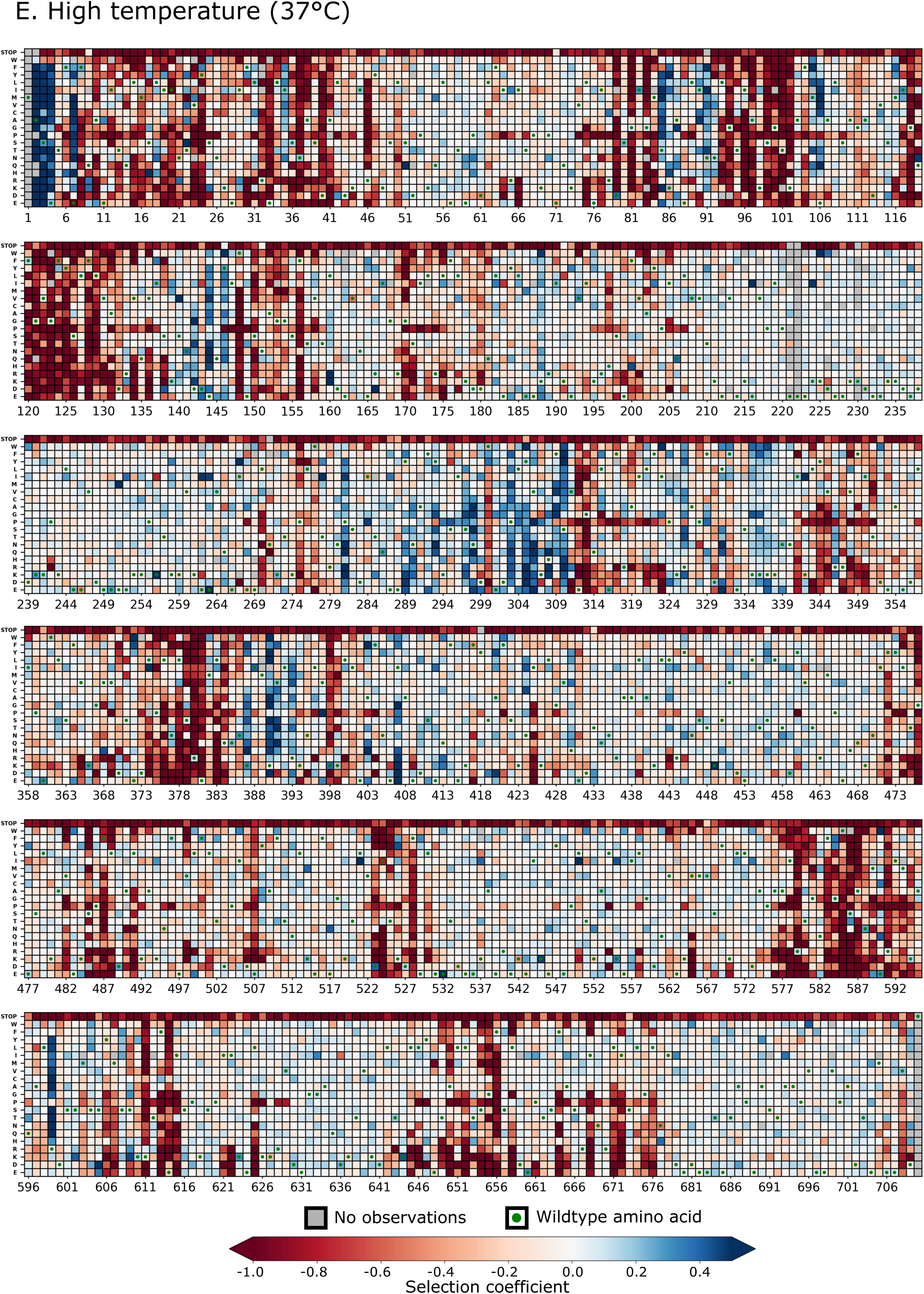
Heatmap representation of the fitness map observed for single amino acid changes across amino acids 2-709 of Hsp90 in (A) nitrogen depletion, (B) high salt, (C) ethanol stress, (D) diamide, and (E) high temperature (37°C).

**Supplementary Figure 2.**
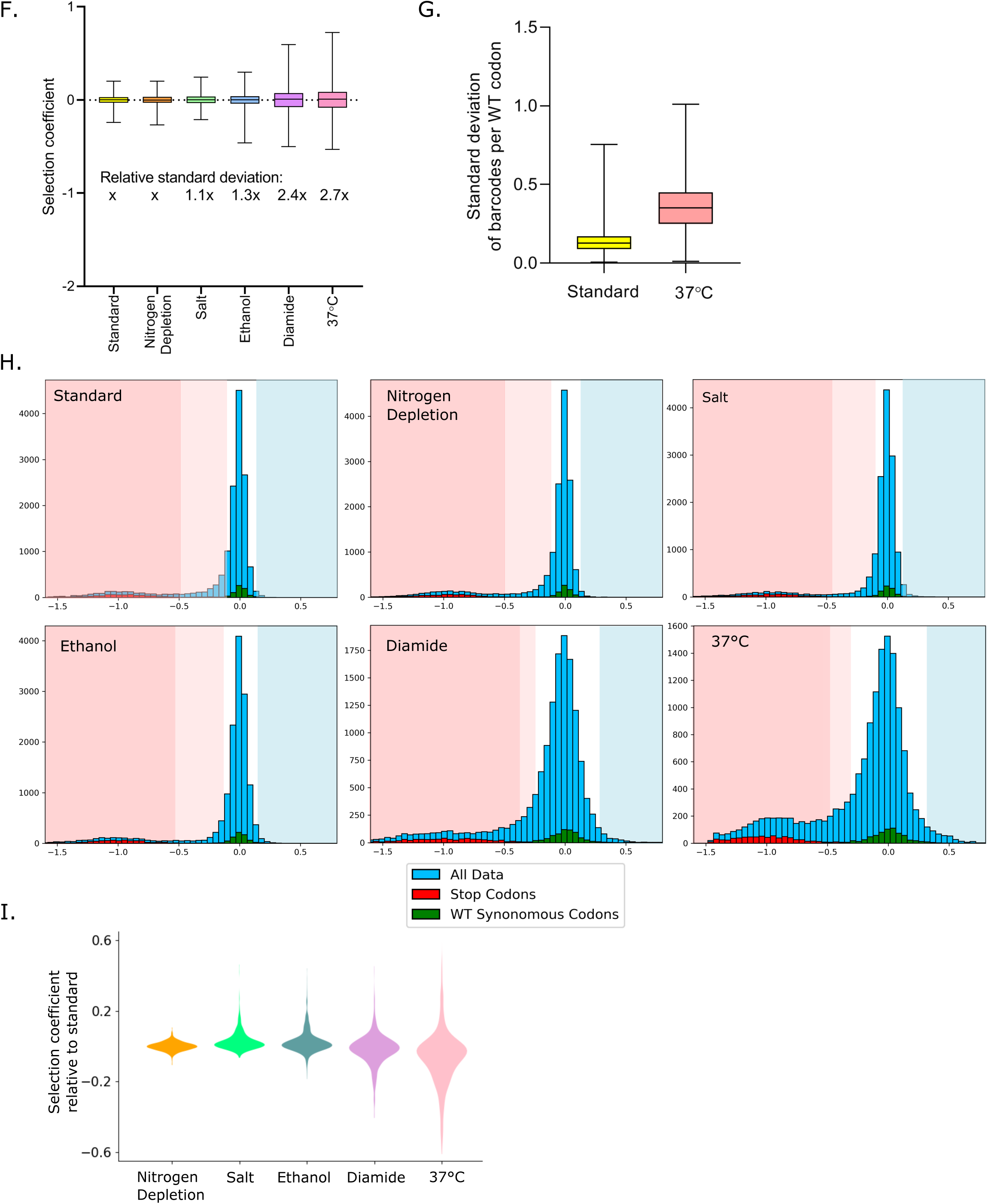
F. The distribution of selection coeffcients for the wildtype synonyms had greater variation in diamide and 37°C than in standard conditions. The standard of deviation of wildtype synonyms in each condition relative to standard conditions (x) is specified under the corresponding box. G. The variation of barcodes for each wildtype codon was higher at 37°C compared to standard conditions. H. Distribution of selection coeffcients in each environmental condition. Mutations were categorized as beneficial (light blue shading), wildtype-like (white shading), intermediate (light pink shading) or deleterious (dark pink shading) based on the distribution of wildtype synonyms (green bars) and stop codons (red bars) in each condition. I. Distribution of the difference between selection coeffcients of each mutation in each stress condition compared to the same mutation in standard conditions.

**Supplementary Figure 3.**
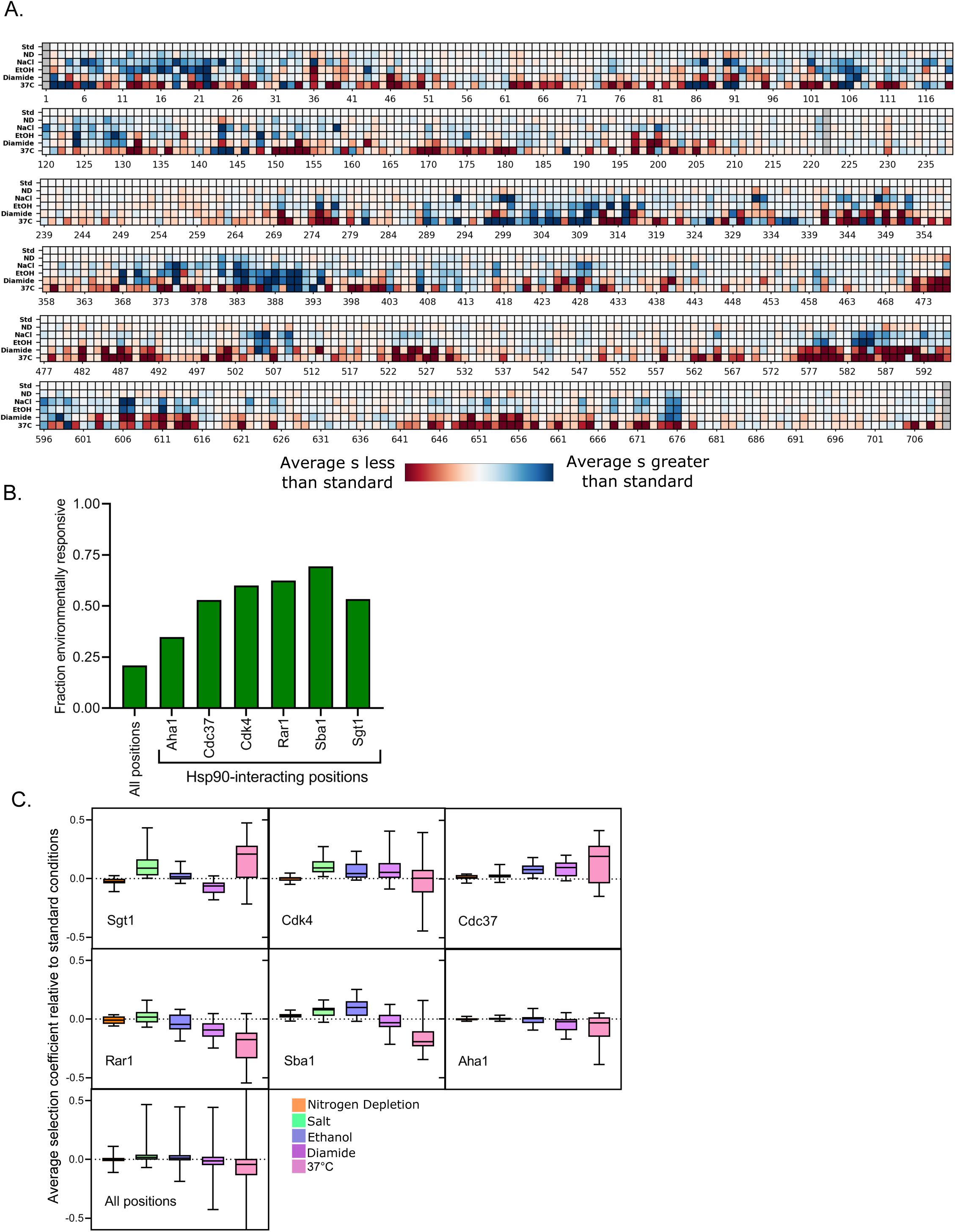
A. Heatmap representation of the average selection coeffcient (excluding stops and wildtype synonyms) at each position in each environmental condition relative to the average standard selection coeffcient at the same position in standard conditions. B. The fraction of Hsp90 positions at interfaces that were categorized as environmentally responsive. C. The average selection coeffcient in each environment relative to standard at all the Hsp90 positions at each stated interface.

**Supplementary Figure 5.**
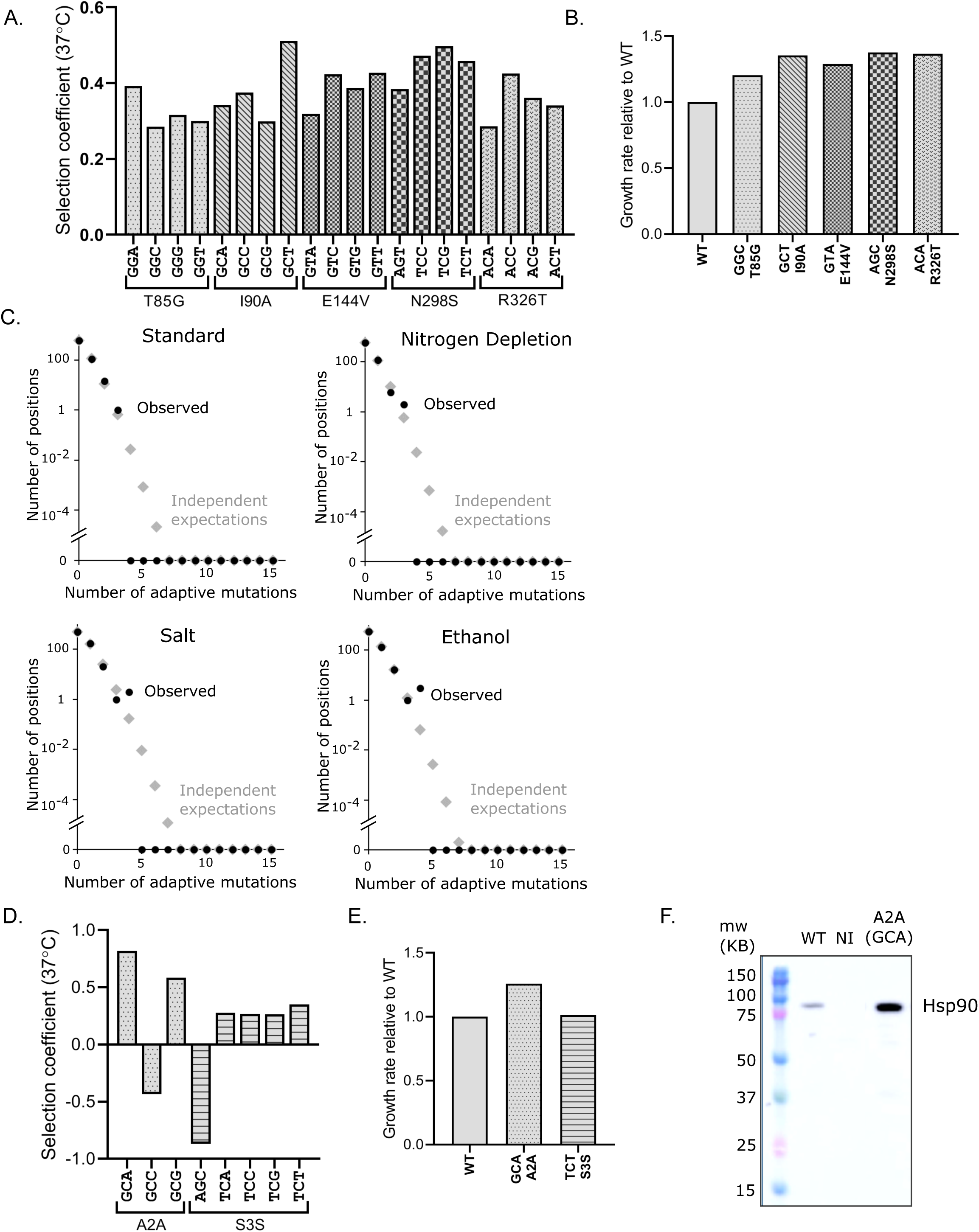
A. Selection coeffcients of synonymous codon variants for five amino acid mutants with beneficial selection coeffcients at 37°C show high correlation. B. The same individual variants analyzed in isolation exhibit increased growth rates. C. Distribution of the number of beneficial mutations at the same position in standard, nitrogen depletion, high salt, and ethanol conditions. D. Selection coeffcients for codon variants of synonymous mutants at the very N-terminus of Hsp90 show high variation. E. Individual synonymous mutant variants with beneficial selection coeffcients exhibited increased growth rates. F. Individual synonymous mutant variants with beneficial selection coeffcients exhibited higher cellular expression levels.

## REFERENCES

1. Ali MM, Roe SM, Vaughan CK, Meyer P, Panaretou B, Piper PW, Prodromou C, Pearl LH (2006) Crystal structure of an Hsp90-nucleotide-p23/Sba1 closed chaperone complex. Nature 440: 1013–1017

2. Bank C, Hietpas RT, Wong A, Bolon DN, Jensen JD (2014) A bayesian MCMC approach to assess the complete distribution of fitness effects of new mutations: uncovering the potential for adaptive walks in challenging environments. Genetics 196: 841–852

3. Bershtein S, Mu W, Serohijos AW, Zhou J, Shakhnovich EI (2013) Protein quality control acts on folding intermediates to shape the effects of mutations on organismal fitness. Molecular cell 49: 133–144

4. Bohen SP, Yamamoto KR (1993) Isolation of Hsp90 mutants by screening for decreased steroid receptor function. Proc Natl Acad Sci U S A 90: 11424–11428

5. Borkovich KA, Farrelly FW, Finkelstein DB, Taulien J, Lindquist S (1989) hsp82 is an essential protein that is required in higher concentrations for growth of cells at higher temperatures. Mol Cell Biol 9: 3919–3930

6. Boucher JI, Bolon DN, Tawfik DS (2016) Quantifying and understanding the fitness effects of protein mutations: Laboratory versus nature. Protein Sci 25: 1219–1226

7. Boucher JI, Cote P, Flynn J, Jiang L, Laban A, Mishra P, Roscoe BP, Bolon DN (2014) Viewing protein fitness landscapes through a next-gen lens. Genetics 198: 461–471

8. Canale AS, Cote-Hammarlof PA, Flynn JM, Bolon DN (2018) Evolutionary mechanisms studied through protein fitness landscapes. Current opinion in structural biology 48: 141–148

9. Chakravarty S, Varadarajan R (1999) Residue depth: a novel parameter for the analysis of protein structure and stability. Structure 7: 723–732

10. Cowen LE, Lindquist S (2005) Hsp90 potentiates the rapid evolution of new traits: drug resistance in diverse fungi. Science 309: 2185–2189

11. Dandage R, Pandey R, Jayaraj G, Rai M, Berger D, Chakraborty K (2018) Differential strengths of molecular determinants guide environment specific mutational fates. PLoS genetics 14: e1007419

12. Darwin C (1859) On the origin of species by means of natural selection, London: John Murray.

13. Darwin C, Wallace A (1858) On the tendency of species to form varieties; and on the perpetuation of varieties and species by natural means of selection. Journal of hte Proceedings of the Linnean Society of London Zoology: 45–62

14. Dingens AS, Arenz D, Overbaugh J, Bloom JD (2019) Massively Parallel Profiling of HIV-1 Resistance to the Fusion Inhibitor Enfuvirtide. Viruses 11

15. Doud MB, Lee JM, Bloom JD (2018) How single mutations affect viral escape from broad and narrow antibodies to H1 influenza hemagglutinin. Nature communications 9: 1386

16. Dykhuizen DE, Dean AM, Hartl DL (1987) Metabolic flux and fitness. Genetics 115: 25–31

17. Firnberg E, Labonte JW, Gray JJ, Ostermeier M (2014) A comprehensive, high-resolution map of a gene’s fitness landscape. Molecular biology and evolution 31: 1581–1592

18. Fisher RA (1930) The Genetical Theory of Natural Selection. Clarendon Press, Oxford

19. Fowler DM, Araya CL, Fleishman SJ, Kellogg EH, Stephany JJ, Baker D, Fields S (2010) High-resolution mapping of protein sequence-function relationships. Nature methods 7: 741–746

20. Fragata I, Blanckaert A, Dias Louro MA, Liberles DA, Bank C (2019) Evolution in the light of fitness landscape theory. Trends in ecology & evolution 34: 69–82

21. Fragata I, Matuszewski S, Schmitz MA, Bataillon T, Jensen JD, Bank C (2018) The fitness landscape of the codon space across environments. Heredity 121: 422–437

22. Gasch AP, Spellman PT, Kao CM, Carmel-Harel O, Eisen MB, Storz G, Botstein D, Brown PO (2000) Genomic expression programs in the response of yeast cells to environmental changes. Molecular biology of the cell 11: 4241–4257

23. Genest O, Reidy M, Street TO, Hoskins JR, Camberg JL, Agard DA, Masison DC, Wickner S (2013) Uncovering a region of heat shock protein 90 important for client binding in E. coli and chaperone function in yeast. Mol Cell 49: 464–473

24. Hagn F, Lagleder S, Retzlaff M, Rohrberg J, Demmer O, Richter K, Buchner J, Kessler H (2011) Structural analysis of the interaction between Hsp90 and the tumor suppressor protein p53. Nat Struct Mol Biol 18: 1086–1093

25. Hawle P, Siepmann M, Harst A, Siderius M, Reusch HP, Obermann WM (2006) The middle domain of Hsp90 acts as a discriminator between different types of client proteins. Mol Cell Biol 26: 8385–8395

26. Hietpas R, Roscoe B, Jiang L, Bolon DN (2012) Fitness analyses of all possible point mutations for regions of genes in yeast. Nature protocols 7: 1382–1396

27. Hietpas RT, Bank C, Jensen JD, Bolon DNA (2013) Shifting fitness landscapes in response to altered environments. Evolution 67: 3512–3522

28. Hietpas RT, Bank, C., Jensen, J.D and Bolon, D. N. (2013) Shifting Fitness Landscapes In Response To Altered Enviornments. Evolution.

29. Hietpas RT, Jensen JD, Bolon DN (2011) Experimental illumination of a fitness landscape. Proc Natl Acad Sci U S A 108: 7896–7901

30. Jarosz DF, Taipale M, Lindquist S (2010) Protein homeostasis and the phenotypic manifestation of genetic diversity: principles and mechanisms. Annual review of genetics 44: 189–216

31. Jiang L, Liu P, Bank C, Renzette N, Prachanronarong K, Yilmaz LS, Caffrey DR, Zeldovich KB, Schiffer CA, Kowalik TF, Jensen JD, Finberg RW, Wang JP, Bolon DNA (2016) A Balance between Inhibitor Binding and Substrate Processing Confers Influenza Drug Resistance. J Mol Biol 428: 538–553

32. Jiang L, Mishra P, Hietpas RT, Zeldovich KB, Bolon DN (2013) Latent effects of hsp90 mutants revealed at reduced expression levels. PLoS Genet 9: e1003600

33. Kacser H, Burns JA (1981) The molecular basis of dominance. Genetics 97: 639–666

34. Kemble H, Nghe P, Tenaillon O (2019) Recent insights into the genotype-phenotype relationship from massively parallel genetic assays. Evolutionary applications 12: 1721–1742

35. Kravats AN, Hoskins JR, Reidy M, Johnson JL, Doyle SM, Genest O, Masison DC, Wickner S (2018) Functional and physical interaction between yeast Hsp90 and Hsp70. Proc Natl Acad Sci U S A 115: E2210–E2219

36. Krukenberg KA, Street TO, Lavery LA, Agard DA (2011) Conformational dynamics of the molecular chaperone Hsp90. Q Rev Biophys 44: 229–255

37. Lang GI, Desai MM (2014) The spectrum of adaptive mutations in experimental evolution. Genomics 104: 412–416

38. Li GW (2015) How do bacteria tune translation efficiency? Current opinion in microbiology 24: 66–71

39. Lindquist S (1981) Regulation of protein synthesis during heat shock. Nature 293: 311–314

40. Lorenz OR, Freiburger L, Rutz DA, Krause M, Zierer BK, Alvira S, Cuellar J, Valpuesta JM, Madl T, Sattler M, Buchner J (2014) Modulation of the Hsp90 chaperone cycle by a stringent client protein. Mol Cell 53: 941–953

41. Lynch M (2013) Evolutionary diversification of the multimeric states of proteins. Proceedings of the National Academy of Sciences of the United States of America 110: E2821–2828

42. Martin G, Lenormand T (2006) A general multivariate extension of Fisher’s geometrical model and the distribution of mutation fitness effects across species. Evolution; international journal of organic evolution 60: 893–907

43. Mavor D, Barlow K, Thompson S, Barad BA, Bonny AR, Cario CL, Gaskins G, Liu Z, Deming L, Axen SD, Caceres E, Chen W, Cuesta A, Gate RE, Green EM, Hulce KR, Ji W, Kenner LR, Mensa B, Morinishi LS, Moss SM, Mravic M, Muir RK, Niekamp S, Nnadi CI, Palovcak E, Poss EM, Ross TD, Salcedo EC, See SK, Subramaniam M, Wong AW, Li J, Thorn KS, Conchuir SO, Roscoe BP, Chow ED, DeRisi JL, Kortemme T, Bolon DN, Fraser JS (2016) Determination of ubiquitin fitness landscapes under different chemical stresses in a classroom setting. eLife 5

44. McCarthy MI, Abecasis GR, Cardon LR, Goldstein DB, Little J, Ioannidis JP, Hirschhorn JN (2008) Genome-wide association studies for complex traits: consensus, uncertainty and challenges. Nature reviews Genetics 9: 356–369

45. Meyer P, Prodromou C, Hu B, Vaughan C, Roe SM, Panaretou B, Piper PW, Pearl LH (2003) Structural and functional analysis of the middle segment of hsp90: implications for ATP hydrolysis and client protein and cochaperone interactions. Mol Cell 11: 647–658

46. Meyer P, Prodromou C, Liao C, Hu B, Roe SM, Vaughan CK, Vlasic I, Panaretou B, Piper PW, Pearl LH (2004) Structural basis for recruitment of the ATPase activator Aha1 to the Hsp90 chaperone machinery. EMBO J 23: 1402–1410

47. Mishra P, Flynn JM, Starr TN, Bolon DNA (2016) Systematic Mutant Analyses Elucidate General and Client-Specific Aspects of Hsp90 Function. Cell reports 15: 588–598

48. Mustonen V, Lassig M (2009) From fitness landscapes to seascapes: non-equilibrium dynamics of selection and adaptation. Trends in genetics : TIG 25: 111–119

49. Nathan DF, Lindquist S (1995) Mutational analysis of Hsp90 function: interactions with a steroid receptor and a protein kinase. Mol Cell Biol 15: 3917–3925

50. Ohta T (1973) Slightly deleterious mutant substitutions in evolution. Nature 246: 96–98

51. Picard D, Khursheed B, Garabedian MJ, Fortin MG, Lindquist S, Yamamoto KR (1990) Reduced levels of hsp90 compromise steroid receptor action in vivo. Nature 348: 166–168

52. Piper PW (1995) The heat shock and ethanol stress responses of yeast exhibit extensive similarity and functional overlap. FEMS microbiology letters 134: 121–127

53. Plotkin JB, Kudla G (2011) Synonymous but not the same: the causes and consequences of codon bias. Nature reviews Genetics 12: 32–42

54. Queitsch C, Sangster TA, Lindquist S (2002) Hsp90 as a capacitor of phenotypic variation. Nature 417: 618–624

55. Retzlaff M, Stahl M, Eberl HC, Lagleder S, Beck J, Kessler H, Buchner J (2009) Hsp90 is regulated by a switch point in the C-terminal domain. EMBO reports 10: 1147–1153

56. Roe SM, Ali MM, Meyer P, Vaughan CK, Panaretou B, Piper PW, Prodromou C, Pearl LH (2004) The Mechanism of Hsp90 regulation by the protein kinase-specific cochaperone p50(cdc37). Cell 116: 87–98

57. Rohl A, Rohrberg J, Buchner J (2013) The chaperone Hsp90: changing partners for demanding clients. Trends Biochem Sci 38: 253–262

58. Rohner N, Jarosz DF, Kowalko JE, Yoshizawa M, Jeffery WR, Borowsky RL, Lindquist S, Tabin CJ (2013) Cryptic variation in morphological evolution: HSP90 as a capacitor for loss of eyes in cavefish. Science 342: 1372–1375

59. Rutherford SL, Lindquist S (1998) Hsp90 as a capacitor for morphological evolution. Nature 396: 336–342

60. Starr TN, Flynn JM, Mishra P, Bolon DNA, Thornton JW (2018) Pervasive contingency and entrenchment in a billion years of Hsp90 evolution. Proceedings of the National Academy of Sciences of the United States of America 115: 4453–4458

61. Stiffler MA, Hekstra DR, Ranganathan R (2015) Evolvability as a function of purifying selection in TEM-1 beta-lactamase. Cell 160: 882–892

62. Taipale M, Krykbaeva I, Koeva M, Kayatekin C, Westover KD, Karras GI, Lindquist S (2012) Quantitative analysis of HSP90-client interactions reveals principles of substrate recognition. Cell 150: 987–1001

63. Tuller T, Carmi A, Vestsigian K, Navon S, Dorfan Y, Zaborske J, Pan T, Dahan O, Furman I, Pilpel Y (2010) An evolutionarily conserved mechanism for controlling the efficiency of protein translation. Cell 141: 344–354

64. Tutt JW (1896) British moths, London: Routledge.

65. Verba KA, Wang RY, Arakawa A, Liu Y, Shirouzu M, Yokoyama S, Agard DA (2016) Atomic structure of Hsp90-Cdc37-Cdk4 reveals that Hsp90 traps and stabilizes an unfolded kinase. Science 352: 1542–1547

66. Wayne N, Bolon DN (2007) Dimerization of Hsp90 is required for in vivo function. Design and analysis of monomers and dimers. J Biol Chem 282: 35386–35395

67. Wright S (1932) The roles of mutation, inbreeding, crossbreeding, and selection in evolution. Proceedings of the Sixth International Congress on Genetics: 355–366

68. Zhang M, Kadota Y, Prodromou C, Shirasu K, Pearl LH (2010) Structural basis for assembly of Hsp90-Sgt1-CHORD protein complexes: implications for chaperoning of NLR innate immunity receptors. Mol Cell 39: 269–281

